# Genome-wide screens for mitonuclear co-regulators uncover links between compartmentalized metabolism and mitochondrial gene expression

**DOI:** 10.1101/2023.02.11.528118

**Authors:** Nicholas J. Kramer, Gyan Prakash, Karine Choquet, Iliana Soto, Boryana Petrova, Hope E. Merens, Naama Kanarek, L. Stirling Churchman

## Abstract

Mitochondrial oxidative phosphorylation (OXPHOS) complexes are assembled from proteins encoded by both nuclear and mitochondrial DNA. These dual-origin enzymes pose a complex gene regulatory challenge for cells, in which gene expression must be coordinated across organelles using distinct pools of ribosomes. How cells produce and maintain the accurate subunit stoichiometries for these OXPHOS complexes remains largely unknown. To identify genes involved in dual-origin protein complex synthesis, we performed FACS-based genome-wide screens analyzing mutant cells with unbalanced levels of mitochondrial- and nuclear-encoded subunits of cytochrome *c* oxidase (Complex IV). We identified novel genes involved in OXPHOS biogenesis, including two uncharacterized genes: *PREPL* and *NME6*. We found that PREPL specifically regulates Complex IV biogenesis by interacting with mitochondrial protein synthesis machinery, while NME6, an uncharacterized nucleoside diphosphate kinase (NDPK), controls OXPHOS complex biogenesis through multiple mechanisms reliant on its NDPK domain. First, NME6 maintains local mitochondrial pyrimidine triphosphate levels essential for mitochondrial RNA abundance. Second, through stabilizing interactions with RCC1L, NME6 modulates the activity of mitoribosome regulatory complexes, leading to disruptions in mitoribosome assembly and mitochondrial RNA pseudouridylation. Taken together, we propose that NME6 acts as a link between compartmentalized mitochondrial metabolites and mitochondrial gene expression. Finally, we present these screens as a resource, providing a catalog of genes involved in mitonuclear gene regulation and OXPHOS biogenesis.

## Introduction

In order to build a healthy mitochondrial proteome, cells are challenged with coordinating gene expression programs across two distinct and physically separated genomes^1–3^. Mitochondria maintain their own extrachromosomal genome, which in humans encodes 13 subunits of oxidative phosphorylation (OXPHOS) complexes – enzymes responsible for cellular energy conversion and aerobic life. The remainder of the OXPHOS complex subunits are encoded in the nucleus, synthesized in the cytosol, and imported into mitochondria for proper OXPHOS complex assembly. This gene regulatory challenge is critical for the cell to overcome as dysfunctional mitochondria are implicated in a variety of human diseases, ranging from neurodegeneration to cancer^4–8^. Despite their importance to human health, a mechanistic understanding of mitochondrial gene expression regulation remains well behind that of its nuclear counterpart.

At every stage of mitochondrial gene expression, from mtDNA replication to transcription and translation, the mitochondrial genome is controlled by nuclear-encoded factors^3^. While the suite of known proteins controlling mitochondrial gene expression has expanded in recent years (for review, see^9,10^), a mechanistic understanding of these regulatory pathways, which likely contribute to the co-regulated expression of nuclear and mitochondrial-encoded OXPHOS subunits, remains unknown. Nuclear- and mitochondrial-encoded OXPHOS subunits are synthesized in a remarkably balanced manner across a variety of human cell lines^11^; however, how a cell achieves this state of mitonuclear balance is poorly understood. Failure to synthesize the correct stoichiometries of OXPHOS subunits may result in a state of ‘mitonuclear imbalance’ leading to cellular stress responses, such as the mitochondrial unfolded protein response, oxidative and metabolic stress responses^12–15^. Moreover, mitonuclear imbalance caused by disruptions in mitoribosome subunit levels has been implicated in longevity and aging phenotypes in yeast, worms, and mice^16–19^. Together, these studies suggest the establishment and maintenance of a healthy mitochondrial proteome via proper mito-cellular communication are critical for cellular physiology.

To identify regulators of mitonuclear balance, we performed FACS-based genome-wide CRISPR screens to uncover factors that altered the accumulation of two dual-genome encoded OXPHOS subunits of Complex IV – COX1 and COX4. Our screens found known regulators of OXPHOS gene regulation and assembly, as well as novel genes with uncharacterized mitochondrial functions whose depletion led to imbalanced OXPHOS subunit levels. Here, we investigated two poorly characterized genes, *PREPL* and *NME6*, both of which altered Complex IV subunit accumulation but in opposite directions. We found that *PREPL* encodes for a dual cytosolic- and mitochondrial-localized protein that is enriched in the nervous system and interacts with mitochondrial protein synthesis machinery to specifically regulate the biogenesis of OXPHOS Complex IV. Furthermore, we observed that NME6, a member of the nucleotide diphosphate kinase (NDPK) family, enzymes which catalyze the transfer of phosphate moieties from nucleoside diphosphates (NDPs) to nucleotide triphosphates (NTPs),^20–22^ controlled OXPHOS biogenesis through multifaceted mechanisms dependent on its NDPK domain. We found that NME6 establishes mitochondrial pyrimidine triphosphate pools essential for maintaining mitochondrial (mt)-encoded RNA abundance. Moreover, through stabilizing interactions with RCC1L, NME6 modulates the activity of mitoribosome regulatory factors and mt-RNA pseudouridylation. Thus, NME6 is a key enzyme connecting compartmentalized mitochondrial metabolite regulation^23^ to mitochondrial gene expression. These data lead to a model where NME6/RCC1L senses the levels of mitochondrial pyrimidines and controls mitochondrial ribosome biogenesis. Altogether, these screens provide a resource of genes for further avenues of research into mito-cellular communication and the regulation of mitochondrial gene expression.

## Results

### FACS-based CRISPR screens for genes regulating mitonuclear balance

In order to uncover genetic regulators of mitonuclear balance, we designed a FACS-based genome-wide screen to identify factors controlling levels of OXPHOS subunits destined for the same complex (Complex IV) but encoded by either nuclear or mitochondrial DNA (**Fig. 1a**). Complex IV assembly, a tightly orchestrated process requiring over 30 assembly factors^24^, begins with the synthesis of mitochondrial-encoded *MT-CO1* (COX1), which must bind nuclear-encoded *COX4I1* (COX4) to initiate assembly^25^. We found that depleting COX4 with sgRNAs targeting *COX4I1* led to both the predicted reduction in COX4 and a concurrent reduction in COX1, consistent with COX1 sensing and responding to levels of COX4, as described previously (**Fig. 1b**)^25^. On the other hand, ethidium bromide or chloramphenicol treatment, inhibitors of mitochondrial transcription and translation, reduced the levels of mitochondrial-encoded COX1 but did not lead to a reduction in nuclear-encoded COX4 (**Fig. 1b,c**). Taken together, these data both established the specificity and dynamic range of our FACS screening conditions and also suggested a directionality to Complex IV biogenesis with mitochondrial gene expression programs poised to respond to nuclear subunit levels.

**Figure 1.**
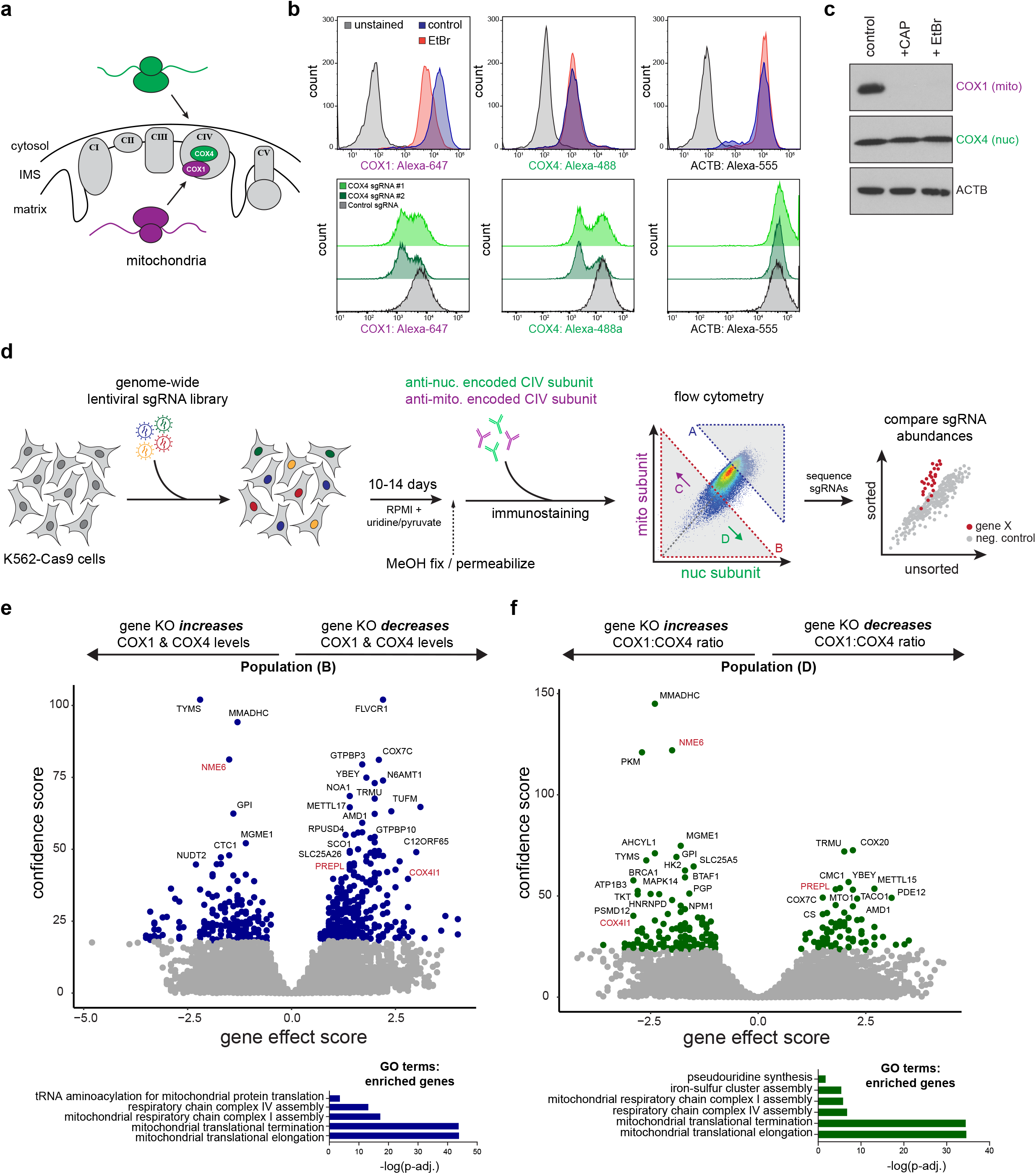
Genome-wide CRISPR screens for mitonuclear balance identify regulators of Complex IV biogenesis. **(a)** Mito-encoded COX1 and nuclear-encoded COX4 are initial assembly subunits of OXPHOS Complex IV. **(b)** FACS immunostaining of COX1, COX4, and ACTB levels after mitochondrial transcriptional inhibition with ethidium bromide (EtBr; 2 µg/mL, 3 days) or with sgRNAs targeting nuclear-encoded *COX4I1*. **(c)** Immunoblotting of mitochondrial and nuclear-encoded subunits after ethidium bromide (2 µg/mL, 5 days) or chloramphenicol treatments (CAP; 200 µg/mL, 5 days). **(d)** Schematic of genome-wide FACS-based CRISPR screening approach to measure mito-nuclear subunit expression. sgRNA libraries include 10 sgRNAs/gene and ∼10,000 negative control sgRNAs. Cells were fixed, immunostained, and sorted based on expression levels of mitochondrial- and nuclear-encoded Complex IV (CIV) subunits (COX1 and COX4, respectively) after 10-14 days. DNA was purified from sorted (Gates A - D) and unsorted control cells to determine sgRNA enrichment or depletion from populations of interest. **(e, f)** Volcano plots summarizing gene knockout (KO) effect (enrichment or depletion of sgRNAs in the sorted population relative to unsorted controls) vs. confidence scores for the sorted populations indicated relative to unsorted control populations (blue or green points = hits < 10% FDR; screens were performed twice, independently). *COX4I1* and hits studied in more detail are shown in red font. Significantly enriched gene ontology terms for positively selected genes (GO biological process, PANTHER) are shown below.

To perform these screens, we measured endogenous levels of mitochondrial (COX1)- and nuclear (COX4)-encoded subunits using FACS in K562-Cas9 cells transduced with a genome-wide single guide RNA (sgRNA) library (10 sgRNAs per gene targeting ∼21,000 human nuclear-encoded genes and ∼10,000 safe-targeting control sgRNAs; **Fig. 1d**)^26^. By analyzing cells with altered levels of both nuclear- and mitochondrial-derived subunits, we disentangled genes essential in mitochondrial health from genes involved in nuclear or mitochondrial gene expression(**Fig. 1e,f, Fig. S1**).

Our screens revealed known players in mitochondrial biology, as well as many novel genetic regulators of OXPHOS biogenesis (**Fig 2a, Table S1**). Positive control sgRNAs targeting *COX4I1* behaved in our screens as predicted, as did genes known to control mitochondrial gene expression at various stages, including mitochondrial transcription (e.g., *TFB2M, POLRMT*), mtRNA processing (e.g., *PDE12, FASTKD5*), mitoribosome subunits and translation factors (e.g., *TUFM, C12ORF65/MTRFR, TACO1*), and Complex IV assembly factors (e.g., *COA3, COA5, COA7, PET117*). However, a number of our top hits were also genes with unknown functions in mitochondrial biology, several of which were predicted to localize to mitochondria in MitoCarta3.0^27^ (**Fig. 2a**).

**Figure 2.**
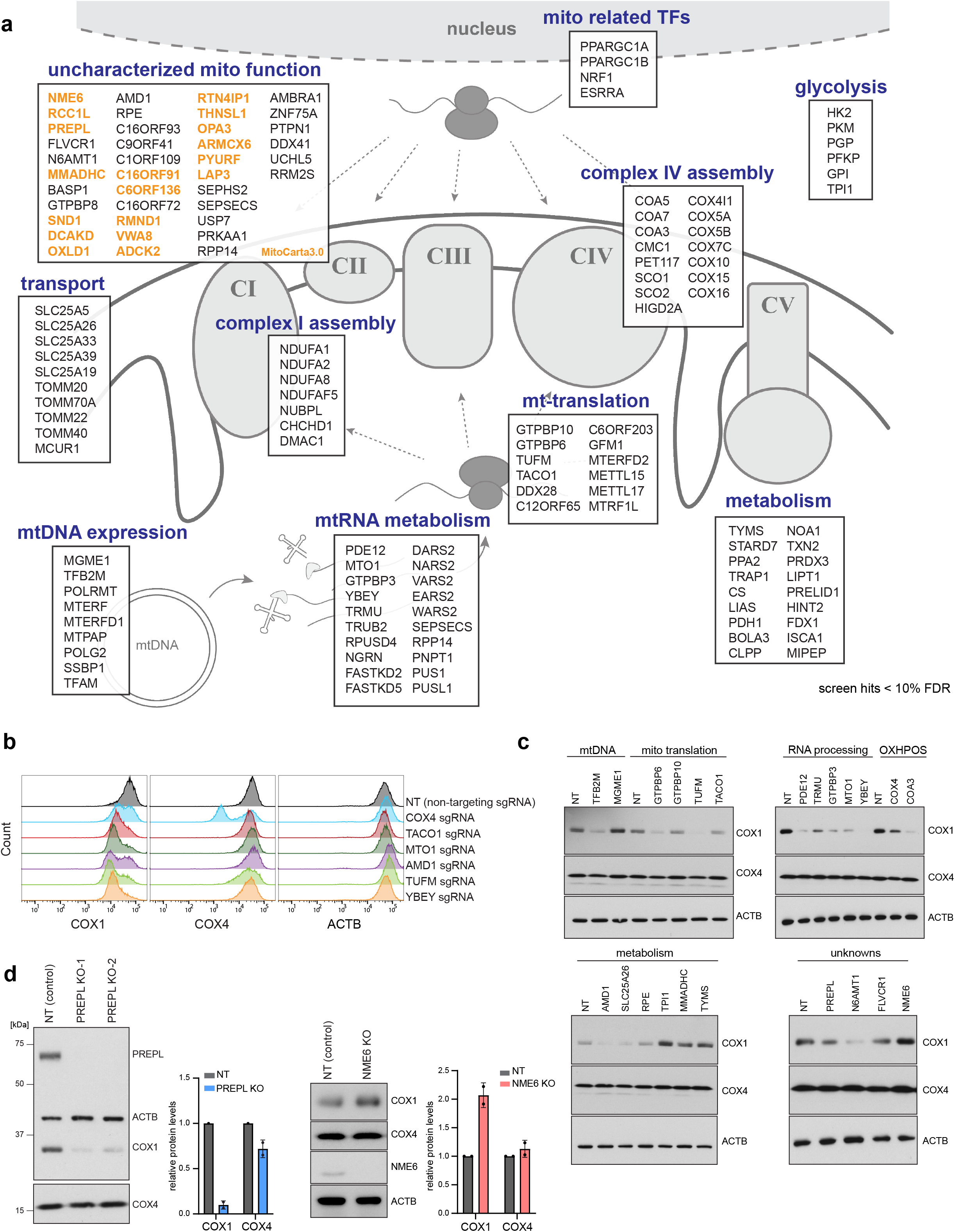
Hit summary of screens for mitonuclear balance. **(a)** Schematic of selected genes (FDR < 10%) grouped by subcellular localization and function. Genes predicted to localize to mitochondria (present in MitoCarta3.0), but with poorly understood functions in mitochondrial biology are highlighted orange (TFs = transcription factors). **(b)** FACS validation of individual sgRNAs in methanol-fixed K562 cells, 10 days post-sgRNA transduction (NT = non-targeting control sgRNA). **(c)** Western blot validation of hits categorized by function, 10 days post-sgRNA transduction. **(d)** Western blot characterization of mitonuclear imbalance phenotypes in homozygous *PREPL* and *NME6* KO K562 cells (2 independently generated clonal KO lines quantified each; normalized to ACTB levels; data are means +/- standard deviation (SD)).

We validated our top hits by infecting cells with individual sgRNAs targeting genes of interest, and subsequently measuring COX1, COX4, and ACTB levels by FACS to rule out non-specific effects on global translation (**Fig. 2b**). Using immunoblots, we then confirmed that several of these hits (∼50 genes tested) altered COX1, COX4, and ACTB expression in the directions predicted from our FACS screens (**Fig 2c, Fig. S2**). Finally, we counter screened these hits by assessing mitochondrial content (MitoTrackerRed), TFAM levels (as a proxy for mtDNA levels), and Complex V subunit levels to disentangle general mitochondrial regulators from Complex IV specific regulators (**Fig. S2**). We further characterized two top hits with uncharacterized mitochondrial functions, which altered COX1/COX4 ratios in opposite directions: *PREPL* (deletion led to a decreased COX1/COX4 ratio) and *NME6* (deletion led to an increased COX1/COX4 ratio) (**Fig. 1e,f**). We generated homozygous single-cell knockout (KO) clones for *PREPL* and *NME6* using CRISPR and confirmed these imbalance phenotypes with immunoblotting (**Fig. 2d**).

### Mitochondrial localized PREPL regulates OXPHOS Complex IV biogenesis

*PREPL* (prolyl-endopeptidase like) is a human disease gene of unknown function, in which mutations cause congenital myasthenic syndrome^28–33^. We found that deletion of *PREPL* in K562 cells caused a reduction in mitochondrial-encoded COX1 while nuclear-encoded COX4 remained near wild-type levels. Reintroduction of cDNA encoding PREPL into *PREPL* KO cells rescued these phenotypes (**Fig. 3a**). *PREPL* is documented in MitoCarta3.0^27^; however, we observed that endogenous PREPL is both cytosolic- and mitochondrial-localized using subcellular cytosolic and mitochondrial fractionation (**Fig. 3b**). We found two dominant PREPL isoforms are expressed in K562 cells – a shorter cytosolic isoform (PREPL_(s)_) and a longer mitochondrial isoform (PREPL_(L)_), consistent with PREPL expression in HEK293T cells^34^. Furthermore, we found a strong tissue specificity in PREPL isoform usage across mouse tissues, as well as a high enrichment of PREPL expression in the brain (**Fig. 3c**).

**Figure 3.**
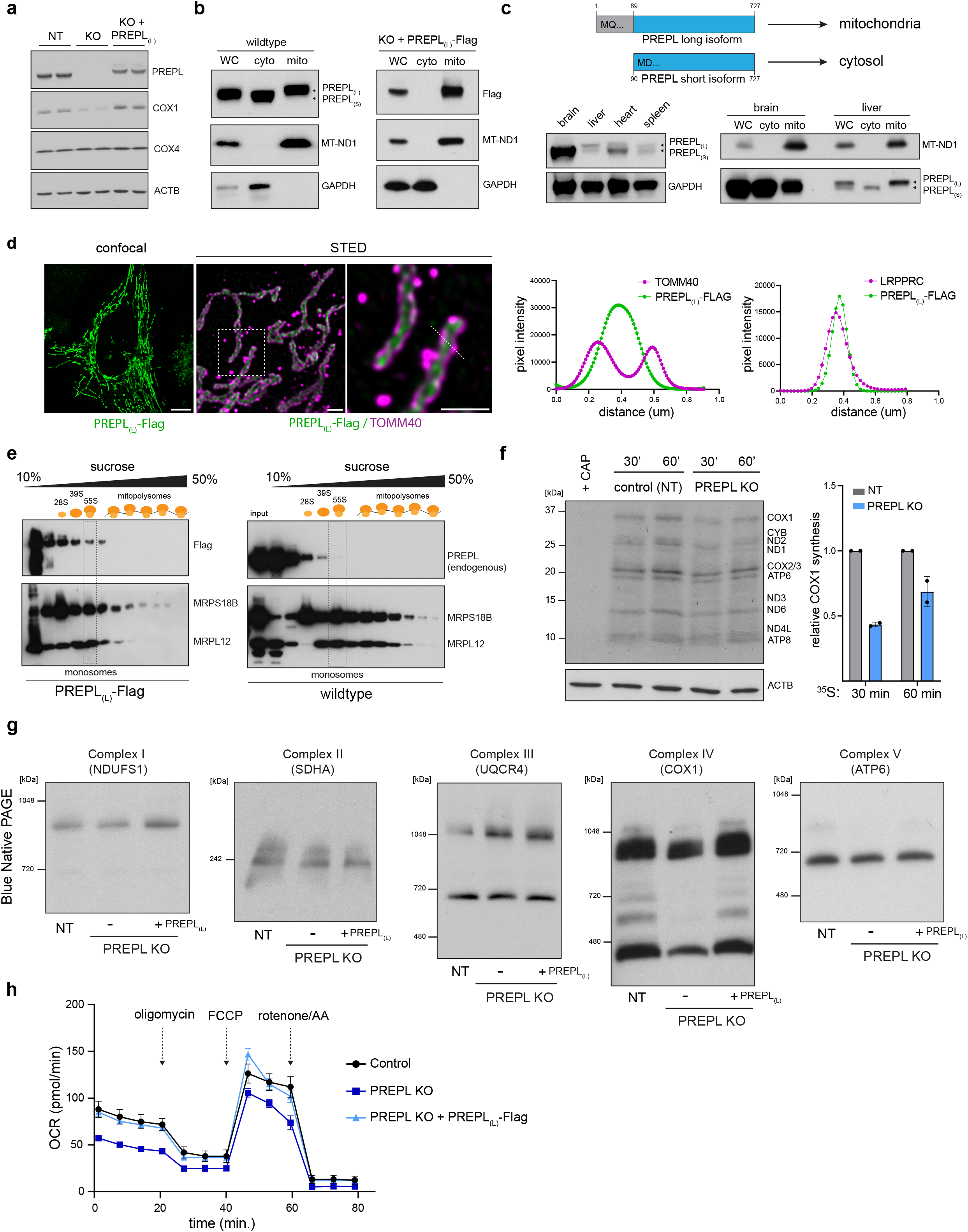
PREPL is a brain-enriched, dual-localized protein regulating OXPHOS Complex IV biogenesis. (**a**) Representative western blots of COX1 and COX4 in NT (non-targeting control sgRNA), PREPL KO, and PREPL KO rescue cells (PREPL-long isoform = PREPL_(L)_). (**b**) Subcellular fractionations and western blotting for endogenous PREPL or exogenous PREPL_(L)_-Flag (WC = whole cell, cyto = cytosol fraction, mito = mitochondrial fraction). (**c**) Schematic representation of short and long PREPL protein isoforms and representative western blot detection of PREPL expression and isoform usage in mouse tissues (wildtype C57BL/6 mice, 8 weeks-old male). (**d**) Left: Representative micrographs of confocal and STED microscopy in U2OS cells transduced with PREPL_(L)_-Flag, stained with anti-Flag and anti-TOMM40 (outer mitochondrial membrane marker) antibodies. Right: representative pixel intensity line scans of fluorescence channels across indicated dashed line (green = FLAG, magenta = TOMM40 or LRPPRC (matrix marker, **Fig S3**); scale bars = 10 µm confocal panel, 500 nm STED panels). (**e**) Western blots of fractions collected from 10-50% sucrose gradients after ultracentrifugation for 3 hours. at 40,000 rpm, 4C (MRPS18B = small mitoribosome subunit, MRPL12 = large mitoribosome subunit). (**f**) Left: ^35^S metabolic labeling for 30 or 60 min. in the presence of anisomycin (100 ug/mL) followed by autoradiography of mitochondrial translation products and western blotting of ACTB to control for loading. Right: Quantification of COX1 synthesis relative to ACTB (CAP = chloramphenicol, 50 ug/mL; n = 2 independent experiments). (**g**) BN-PAGE followed by western blot detection of native OXPHOS Complexes I, II, III, IV, and V. (**h**) Seahorse assay measuring oxygen consumption rate in live cells with mitochondrial stressors (FCCP = carbonyl cyanide-p-trifluoromethoxyphenylhydrazone, AA = antimycin A; n = 4-5 cultures). Data are means +/- SD.

To determine how PREPL controls mitonuclear balance, we identified protein-interactors of PREPL_(L)_ using immunoprecipitation/mass spectrometry. PREPL_(L)_ interactes with many mitochondrial matrix-localized proteins significantly enriched for GO terms including ‘mitochondrial translation’, ‘tRNA metabolic processing, and ‘mitochondrial protein processing’. We confirmed the the interactions with FASTKD4 and LARS2 with co-immunoprecipitations (**Fig. S4, Table S2**). Furthermore, we verified a matrix localization for PREPL_(L)_ using STED superresolution microscopy (**Fig. 3d, S3**).Since PREPL protein interactors GO terms were enriched in mitochondrial translation we tested the PREPL role in mitochondrial translation. Using sucrose gradient ultracentrifugation, we observed that PREPL, as well as FASTKD4, co-sedimented with mitoribosomes, up to the assembled monosome fraction (**Fig. 3e, S4**). Further based on ^35^S-methionine labeling newly synthesized miothcindrial proteins, we found that *PREPL* KO reduced COX1 synthesis without major changes in mtDNA levels, mt-RNA levels, COX1 RNA processing, poly(A) tail length, or COX1 subunit turnover (**Fig. 3f, S4, S5, S6**). These defects led to specific reductions in Complex IV, including the monomeric complex, dimeric, and supercomplex assemblies as determined by blue native PAGE (BN-PAGE) (**Fig. 3g**), and reduction of Complex IV levels were sufficient to cause respiration defects in PREPL KO cells (**Fig. 3h**). Both Complex IV assembly and respiration defects were rescued by expression of mito-PREPL_(L)_ in *PREPL* KO cells. Taken together, our data suggest that PREPL is a brain-enriched, dual cytosolic- and mitochondrial-localized protein that specifically controls Complex IV biogenesis through the regulation of mitochondrial translation.

### NME6 knockout disrupts mitochondrial respiration and OXPHOS biogenesis

Our screens predicted that *NME6* mutation disrupts Complex IV subunit ratios by increasing the levels of mito-encoded COX1 – an opposite effect compared to *PREPL* deletion. NME6, a group II nucleoside diphosphate kinase (NDPK), contains a highly conserved histidine residue within its NDPK domain, residues typically used by NDPK enzymes for phosphate transfer (**Fig. 4a**). While NME6 has no predicted mitochondrial targeting sequence, it is included in MitoCarta3.0^27^. We found that NME6 localized to the mitochondrial matrix using a combination of confocal microscopy, subcellular fractionations, and Proteinase K protection assays in purified mitochondria (**Fig. 4b,c**). NME6 KO cells had reduced proliferation in standard RPMI-1640 media as well as the ‘physiological’ human plasma like media^35^ (HPLM; **Fig. 4d, S7**). These proliferative phenotypes in NME6 KO cells were accompanied by decreased basal and maximal respiration rates (**Fig. 4e**) and dysregulated OXPHOS subunit levels (**Fig. 4f**). To test whether these phenotypes relied on the predicted NDPK function of NME6, we mutated the highly conserved histidine to alanine (H137A). We found that exogenous expression of WT, but not H137A NME6, rescued the proliferation phenotypes and disrupted OXPHOS subunit levels of NME6 KO cells to near WT levels (**Fig. 4d,f**).

**Figure 4.**
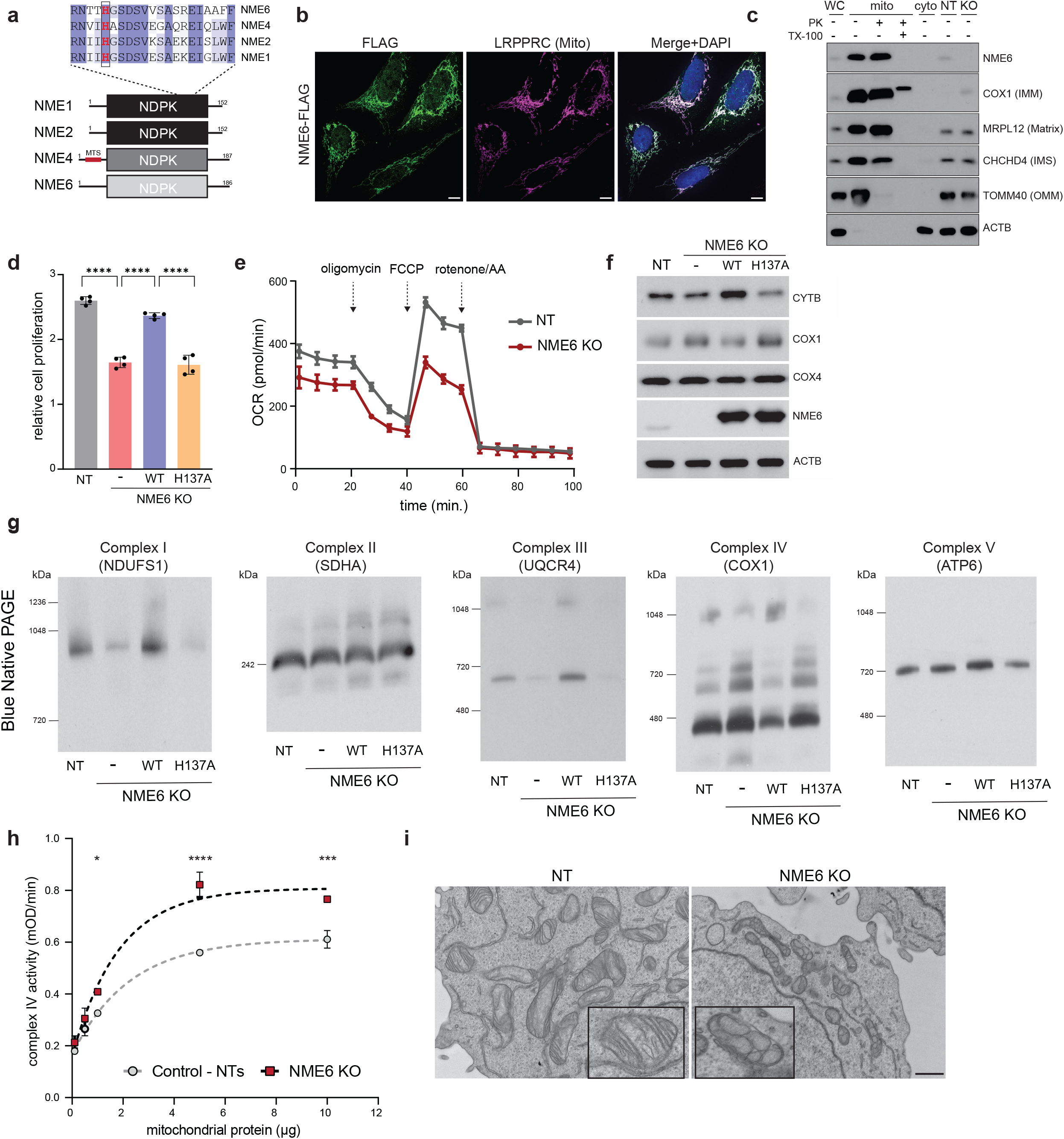
NME6 KO disrupts OXPHOS biogenesis and mitochondrial respiration. (**a**) Schematic of NDPK family members and domain alignment indicating conserved histidine 137 in NME6 (shading represents level of conservation by percent identity using Clustal Omega NME1–NME2 = 88%, NME1–NME4 = 55%, NME1–NME6 = 25%). (**b**) Representative micrographs of U2OS cells transduced with NME6-Flag, immunostained with anti-Flag and anti-LRPPRC antibodies (green = Flag, magenta = LRPPRC, blue = DAPI; scale bar = 10 µm). (**c)** Subcellular fractionations and Proteinase K (PK) digestion of mitochondria in the presence or absence of Triton X-100 (TX-100) detergent followed by western blotting with indicated antibodies (WC = whole cell, cyto = cytosol fraction, mito = mitochondrial fraction, NT= non-targeting sgRNA, KO = NME6 KO). (**d**) Cell proliferation measurements at day 4 relative to day 0 in control (NT) and NME6 KO cells measured by CellTiter-Glo in HPLM media (n = 4, **** = p < 0.0001). (**e**) Seahorse assay assessing oxygen consumption rate (OCR) in live cells using mitochondrial stressors (FCCP = carbonyl cyanide-p-trifluoromethoxyphenylhydrazone, AA = antimycin A; n = 3-4). (**f**) Representative western blots of COX1, COX4 (Complex IV) and CYTB (Complex III) subunit levels in control (NT), KO, and WT/H137A rescue cells. (**g**) BN-PAGE measurements of native OXPHOS complexes in control, NME6 KO, or KO cells expressing WT or H137A NME6. (**h**) Complex IV enzymatic activity in the indicated mitochondrial lysate amounts (n = 2 per protein input, experiments were repeated 2 independent times; * p < 0.05, *** p < 0.001, **** p < 0.0001). (**i**) Representative electron micrographs of mitochondrial cristae structure by transmission electron microscopy in NT and NME6 KO cells (scale bar = 800 nm). Data are means +/- SD.

We next asked what molecular mechanisms could lead to the proliferation and respiration phenotypes we observed in NME6 KO cells. Because NME6 KO cells had elevated levels of COX1 subunits, we tested whether these increased subunits were being functionally integrated into fully assembled complexes and, if so, why increased OXPHOS Complex IV would lead to respiratory defects. Interestingly, using BN-PAGE, we found that NME6 KO had differential effects across OXPHOS complexes; NME6 KO cells had elevated levels of Complex IV but decreased levels of both Complex I and III assemblies (**Fig 4g**). These OXPHOS assembly defects were rescued by reintroducing WT NME6, but not H137A NME6, into NME6 KO cells. We confirmed the upregulation of Complex IV in NME6 KO cells by measuring the enzymatic activity of Complex IV *in vitro* from WT and NME6 KO mitochondrial lysates (**Fig. 4h, S7**). Dysregulation of OXPHOS assembly across complexes in NME6 KO cells was further accompanied by defects in mitochondrial cristae morphology (**Fig. 4i**). Together, NME6 deletion led to disrupted OXPHOS biogenesis and abnormal respiration – phenotypes reliant on a conserved histidine in the NDPK domain of NME6.

### NME6 regulates mitochondrial pyrimidine pools and mt-RNA abundance

Next, we addressed the mechanism by which NME6 acts to regulate mitochondrial gene expression. To test whether mitochondrial mRNA transcripts were affected by NME6 KO, we measured their abundance using a MitoStrings panel^36^, which directly measures RNA abundances with NanoStrings technologies. These data indicated that the majority of mitochondrial mRNAs were downregulated in NME6 KO cells, most severely *MT-CYB*, but with the notable exceptions of *RNR1, RNR2*, and *MT-CO1*, which are more proximal to the heavy strand promoter and were, by contrast, upregulated (**Fig. 5a**). These changes were not concurrent with alterations in mtDNA copy number (**Fig. 5b**). The disruptions in mt-RNA abundances were consistent with the OXPHOS assembly defects we observed where Complex IV biogenesis was upregulated while Complex I and III were downregulated, using BN-PAGE (**Fig. 4g**). Considering the order of polycistronic transcription from mtDNA, these mRNA changes could suggest a defect in transcription elongation or premature termination. NME6 KO did not phenocopy the mutation of the transcription initiation factor TFB2M, which showed near equal reduction across all mitochondrial-encoded transcripts (**Fig. S7**). To directly test the possibility of a transcriptional defect in NME6 KO cells, we pulse-labeled newly synthesized RNAs in purified mitochondria with ^32^P-UTP and visualized newly transcribed RNA with denaturing-PAGE and autoradiography (**Fig. S8**). In these bulk measurements, NME6 KO did not cause large-scale changes in the levels of newly synthesized transcripts, indicating that NME6 KO-mediated changes in RNA levels do not stem from broad transcriptional inhibition.

**Figure 5.**
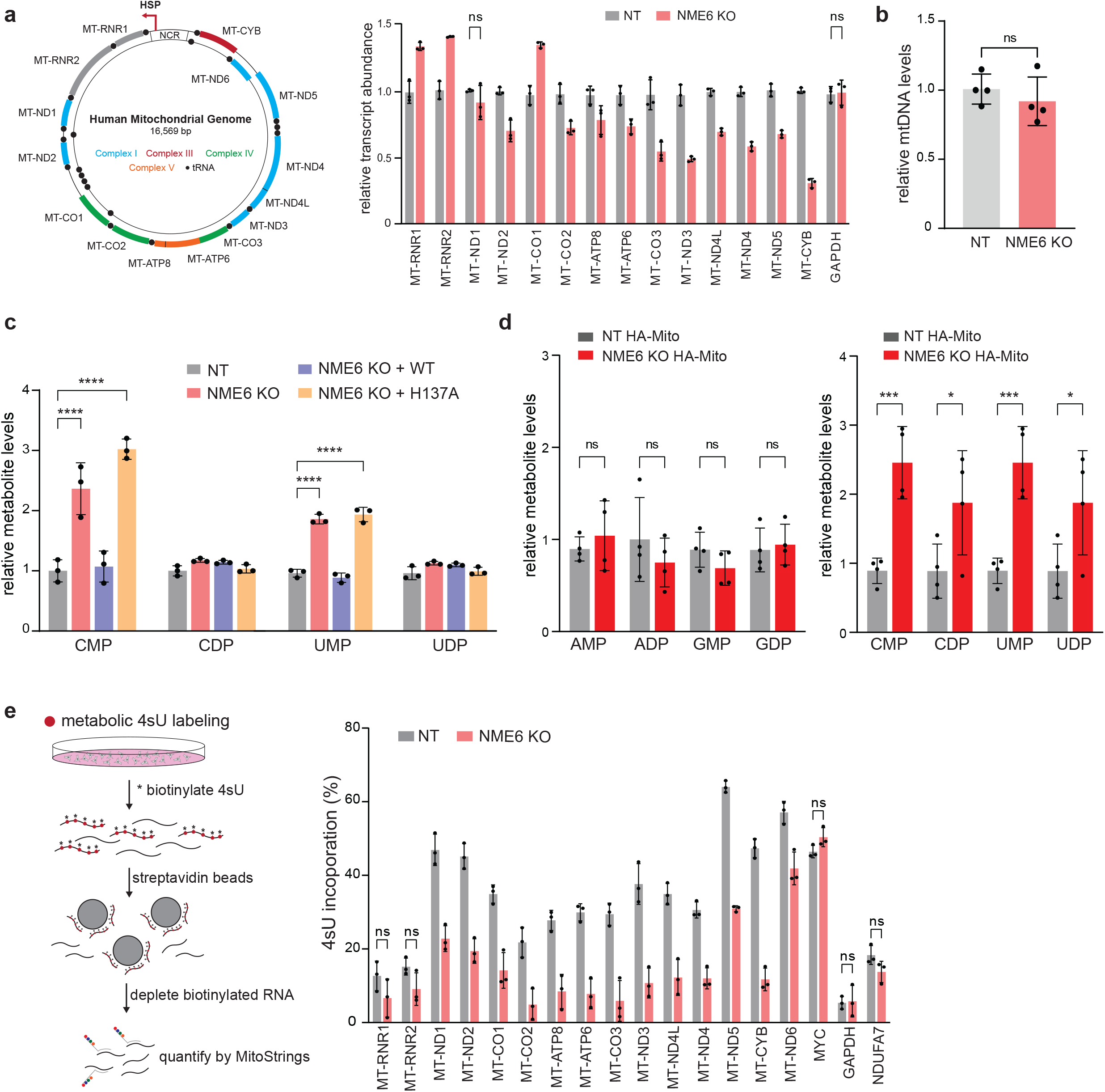
NME6 KO disrupts mitochondrial pyrimidine levels and mt-RNA transcript abundance. (**a**) Left: schematic of human mitochondrial DNA and right: mRNA abundance quantified by MitoStrings, plotted in order of polycistronic transcription (normalized to nuclear-encoded NDUFA7; p < 0.05 unless otherwise stated, n.s. = non-significant, n = 4 replicates from 2 independent experiments). (**b**) qPCR measurements of mtDNA levels normalized to nuclear DNA (mito probe = *MT-TL*, nuclear probe = *B2M;* n = 4 replicates across 2 independent experiments). Mass spectrometry detection of metabolites from (**c**) whole cell or (**d**) mito-IP lysates (normalized to NT for each metabolite; * p < 0.05, *** p < 0.001, **** p < 0.0001; n = 4 independent cultures). (**e**) 4sU transcript incorporation measured by metabolic labeling in control (NT) or NME6 KO cells. Left: schematic of the experimental approach. Right: unlabeled RNAs were quantified by MitoStrings (normalized with a spike-in control; p < 0.05 unless otherwise stated, n.s. = non-significant; n = 3). Data are means +/- SD.

Because our data have demonstrated that the NDPK domain is necessary for NME6 function, we asked whether NME6 was responsible for maintaining local concentrations of mitochondrial nucleotides. Using mass spectrometry-based metabolomics in whole cell lysates, we observed an accumulation of nucleoside monophosphates, including CMP, UMP, and GMP, in NME6 KO cells, suggesting a possible dysregulation of nucleotide biosynthesis; however, only CMP and UMP were dependent on H137A, indicating a role for NME6 specifically in the regulation of pyrimidine levels (**Fig. 5c**). Since whole cell metabolite levels might not be indicative of mitochondria metabolite pools, we analyzed mitochondrial-specific metabolite pools using rapid mitochondrial immunoprecipitation^37^. We found that NME6 KO led to the abnormal accumulation of mono- and diphosphate pyrimidines within mitochondria (**Fig. 5d**) without major disruptions to other general metabolite levels (**Fig. S9**), indicating a disruption in pyrimidine NTP synthesis in NME6 KO mitochondria.

These results suggest that NME6 establishes local pyrimidine NTP levels within mitochondria, which in turn modulates mitochondrial gene expression. Because of the alterations in both steady-state mt-mRNA abundance and pyrimidine levels in NME6 KO cells, we tested the incorporation of pyrimidines into the mitochondrial transcripts. Using 4-thiouridine (4sU) metabolic labeling, followed by 4sU-biotinylation and streptavidin pulldown to deplete newly labeled mt-RNAs, we measured 4sU incorporation by quantifying the amount of unlabeled mt-RNAs using MitoStrings (**Fig. 5e**). NME6 KO reduced the levels of 4sU incorporation for all of the mt-RNAs measured (the highly stable rRNAs did not pass our statistical cut-off), but not for three control nuclear-encoded RNAs. Taken together, these results suggest that NME6 regulates mt-RNA abundance through the establishment of local mitochondrial pyrimidine pools.

### NME6 interacts with RCC1L to control mitoribosome assembly and pseudouridylation levels

NME6 has been reported to interact with the putative guanine exchange factor, RCC1L (WBSCR16), which was also a hit in our screen and is a poorly characterized protein with functions in mitoribosome assembly^38–40^. RCC1L acts through interactions with a larger module of genes known to regulate 16S rRNA stability and RNA pseudouridylation levels, including TRUB2, RPUSD4, and RPUSD3^41,42^. Using IP-mass spectrometry, we confirmed that NME6 and RCC1L interact and found that they bind each other with high specificity (**Fig. 6a, Table S2**). Moreover, in a co-IP analysis, we observed that NME6/RCC1L interactions were reduced in H137A-relative to WT-NME6 and that the stability of either protein depended on the presence of the other (**Fig. 6b, c**).

**Figure 6.**
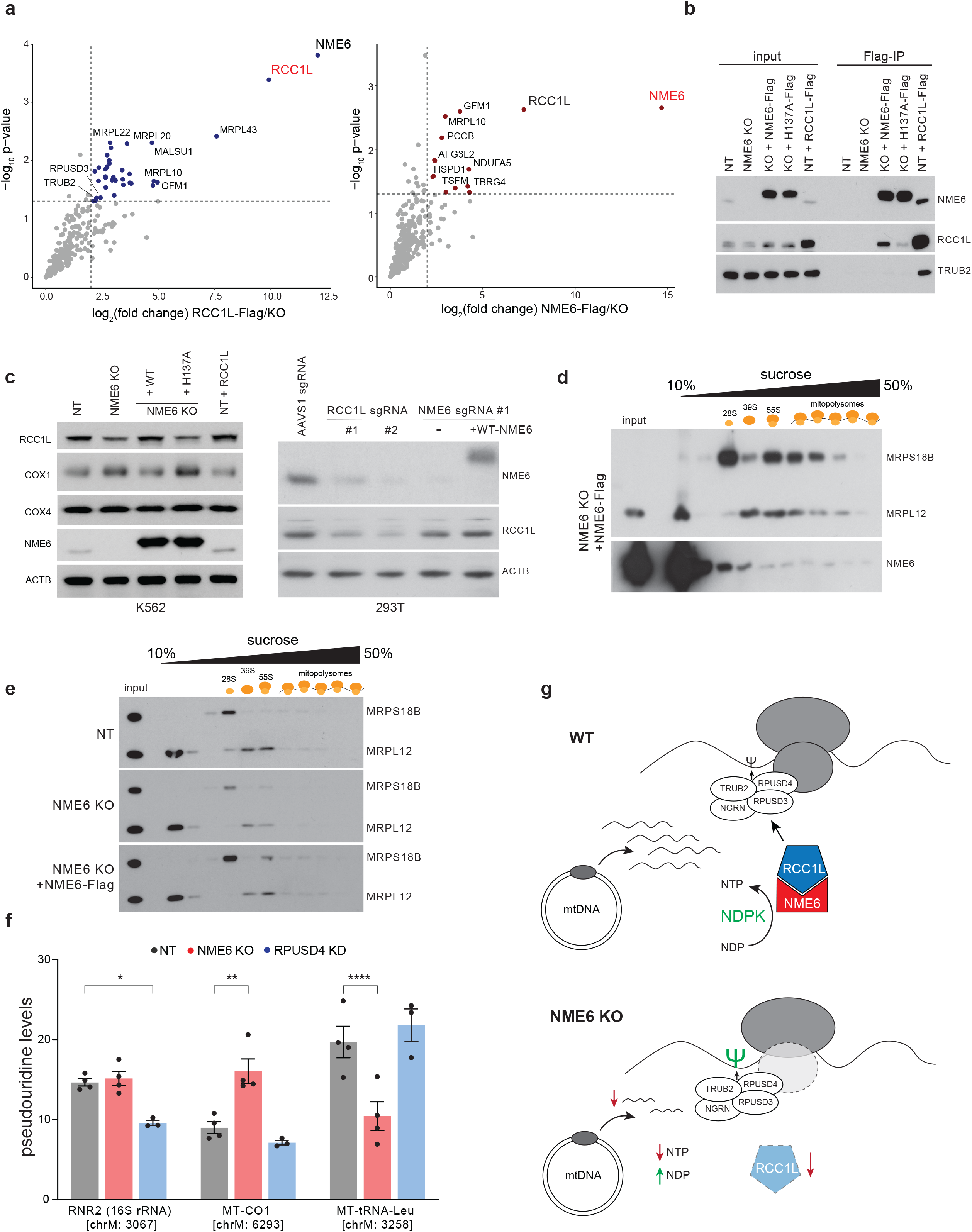
NME6 binds RCC1L to regulate mitoribosome assembly and pseudouridylation levels. (**a**) IP-mass spectrometry identification of proteins interacting with NME6-Flag or RCC1L-Flag in purified mitochondria (colored points = p < 0.05 and log_2_(fold-change) > 2 indicated by dashed lines). (**b**) Co-immunoprecipitation analysis of anti-Flag pulldowns in control (NT), NME6 KO, NME6 KO + WT-NME6-Flag, NME6 KO + H137A-NME6-Flag, or NT + RCC1L-Flag cells. (**c**) Left: western blot analysis of lysates from K562 cell lines (NT, NME6 KO +/- NME6 WT or H137A, and NT+RCC1L) and right: HEK293T cells transduced with indicated sgRNAs (AAVS1 = control sgRNA). (**d** and **e**) Sucrose gradient centrifugation and fractionation followed by western blotting of small (MRPS18B) and large (MRPL12) mitoribosome subunits to monitor (**d**) NME6 co-sedimentation with mitoribosomes and (**e**) mitoribosome assembly phenotypes in NME6 KO cells. (**f**) CMC-sequencing of NT, NME6 KO, or RPUSD4 knockdown (KD) cells to measure pseudouridine levels at three high confidence modification sites in *RNR2, MT-CO1*, and *MT-TL* (data are means +/- SD; * p < 0.05, ** p < 0.01, **** p < 0.0001; n = 3-4 replicates across 2 independent experiments and 2 independently generated clonal KO cell lines). (**g**) Schematic of NME6 function in the regulation of mitochondrial gene expression: 1) local mitochondrial pyrimidine NTP regulation affecting mt-RNA abundance and 2) stability dependent interactions with RCC1L which modulate mitoribosome assembly and the activity of mitoribosome associated pseudouridine synthases (psi = pseudouridine modified base).

Due to this strong co-dependence, we asked whether NME6 has roles in mitoribosome biogenesis and mt-RNA pseudouridylation similar to those reported for RCC1L and the associated pseudouridine synthases^39,42^. We found that NME6 co-sediments with mitoribosomes in sucrose gradients and NME6 KO cells showed assembly defects through disruptions of mainly the small mitoribosome subunit (**Fig. 6d,e, S9**). Using CMC-sequencing, we found that NME6 KO cells had a ∼2-fold increase in pseudouridylation at previously reported modification sites within *MT-CO1* mRNA (chrM: ψ6293), as well as decreased pseudouridine levels within MT-tRNA^Leu^ (chrM: ψ3258) (**Fig. 6f**). NME6 KO did not cause any alteration in the pseudouridine levels in *RNR2* (chrM: ψ3067), a site previously shown to be regulated by the pseudouridine synthase, RPUSD4, which we confirmed here^41^. Together, these data demonstrate a multifaceted role for NME6 in regulating mitochondrial gene expression: NME6 regulates mitochondrial nucleotide levels, mt-RNA abundances, and mitoribosome biogenesis through its interaction with RCC1L.

## Discussion

Here, we performed a series of genome-wide screens using FACS to find gene expression regulators of mitochondrial- and nuclear-encoded subunits of Complex IV. In addition to well-established OXPHOS regulators, among our top-ranked hits were several novel genes with uncharacterized functions in mitochondria. Thus, we believe these screens provide a new high-quality resource for investigating OXPHOS biogenesis. We focus on two such genes here, *PREPL* and *NME6*; however, we uncovered several other genes of interest predicted to regulate OXPHOS biogenesis that will provide interesting avenues for further investigation.

Analysis of mutants with both increased and decreased OXPHOS subunit abundances allowed us to uncover both positive and negative regulators of Complex IV biogenesis. This type of gene regulation based screening is complementary to other screens that have used cell death or growth rate in the presence of OXPHOS inhibitors to uncover OXPHOS regulators^42,43^. In the course of validating hits from our screens, we observed that mitochondrial gene expression appeared quite dynamic, exemplified by our observations that knocking down nuclear-encoded COX4 led to a concurrent decrease in mitochondrial-encoded COX1 to approximately the same extent as the COX4 target itself. Furthermore, we found that most hits we validated altered mitochondrial-encoded COX1 specifically and to a greater extent than nuclear COX4. These data suggest that mitochondrial gene expression is dynamic and prone to multiple sources of regulation, a phenomenon also observed recently in genome-scale Perturb-seq studies^44^, warranting future studies on mitochondrial gene regulation dynamics. Here, we further characterized two such regulators of mitochondrial gene expression, *PREPL* and *NME6*.

*PREPL*, prolyl endopeptidase-like, encodes a putative oligoserine peptidase of unknown function. Mutations in *PREPL* cause congenital myasthenic syndrome 22 (CMS22; OMIM #616224)^29^, an autosomal recessive disorder characterized by neuromuscular transmission defects, neonatal hypotonia, and growth hormone deficiencies. While *PREPL* KO mice exhibit growth and hypotonia phenotypes^45^, the molecular mechanisms of PREPL function remain elusive. We found that within the mitochondrial matrix, PREPL_(L)_ associates with protein synthesis machinery leading to defects in COX1 synthesis. Interestingly, global analysis of mitoribosome associated proteins in mice, found that PREPL interacted with the mitoribosome in a tissue-specific manner^46^. We observed a significant enrichment of PREPL in nervous system tissue, consistent with its genetic involvement in neuromuscular transmission defects in CMS22 patients, but we also observed tissue specific isoform prioritization. Understanding why certain tissues prioritize the expression of one isoform or the other will enable a better understanding of the function of PREPL in both physiology and disease.

Next, we characterized *NME6*, a gene that, when mutated, led to the increased accumulation of mitochondrial-encoded COX1. Our data show that NME6 is mitochondrial localized and plays a key role in regulating mitochondrial gene expression through its conserved NDPK domain. Despite respiratory and proliferation phenotypes, NME6 KO did not simply lead to general reductions in OXPHOS biogenesis. But rather, NME6 is involved in multifaceted OXPHOS gene regulation playing differential roles across each complex.

We found NME6 KO cells had increased levels of the mt-RNAs encoded most proximal to the heavy strand promoter (RNR1, RNR2, MT-CO1) but lower levels of more distal mt-RNAs, e.g. MT-CYB. Importantly, we did not observe gross mitochondrial transcription defects in NME6 KO cells. Nevertheless, our transcriptomics and metabolomics data indicate a role for NME6 in regulating mitochondrial pyrimidine homeostasis and incorporation during transcription via its NDPK domain. However, this role is insufficient to fully explain the differential mt-RNA abundances we observe, particularly for *MT-CO1* mRNA. In fact, we report a second function for NME6 through its interactions with the putative guanine exchange factor, RCC1L.

We found that NME6 binds specifically to RCC1L, which, in turn, interacts with a broader module of genes identified to regulate 16S rRNA stability^42^ and mitoribosome assembly^39^. We observed that the stability of both NME6 and RCC1L rely on the presence of one another, suggesting they exist as obligate heterodimers. Interestingly, RCC1L exhibits guanine exchange factor (GEF) activity^40^, which raises the intriguing possibility that it could bind nucleotides and bring them in proximity to NME6, acting as a potential cofactor for its NDPK activity. RCC1L further interacts with several mitochondrial pseudouridine synthases, including RPUSD4, RPUSD3, and TRUB2^41^. Consistently, we found that NME6 KO cells have increased mt-RNA pseudouridylation, particularly within *MT-CO1* mRNA, a possible explanation for upregulated *MT-CO1* transcript levels or translational output of COX1 in NME6 KO cells. While the function of pseudouridine modifications in human mRNAs is unclear^47,48^, they have been reported to enhance translation efficiency and immune evasion in mRNA-based therapeutics^49–52^.

Together, our data suggest that NME6 has multifaceted functions in mitochondrial gene regulation (**Fig. 6g**), including both the regulation of local mitochondrial pyrimidine nucleotide pools and, together with RCC1L, mitochondrial ribosome assembly, and mt-RNA pseudouridylation. Considering these functions, we propose that NME6/RCC1L acts as a metabolic sensor to tune levels of mitochondrial gene expression based on compartmentalized metabolite availability in the mitochondria. Compartmentalization of metabolites is emerging as a key mechanism underlying important physiological processes, such as proline biosynthesis and mammalian development^23,53–55^. *NME6* is predicted to be an essential gene in mammals based on large-scale mouse KO consortia (IMPC; MGI:1861676) and was also reported as a sensitizing hit for growth in the ‘physiological media’, HPLM^56^. Moreover, *NME6* was found as an essential gene for diffuse midline glioma cancer cell growth, a cancer with a selective dependency on pyrimidine biogenesis^57^. In future work, it will be important to determine which roles of *NME6* contribute to these phenotypes and whether its potential role as a metabolic sensor contributes to interorganellar communication to drive cellular transitions in response to environmental cues.

## Acknowledgements

We’d like to thank R.S. Isaac and C. Guegler for critical reading of the manuscript as well as S. Adamson for help with BN-PAGE experiments. This research was conducted with support from the HMS Electron Microscopy Facility, the HMS Taplin Mass Spectrometry facility, the HMS MicRoN and Neurobiology Imaging Facilities, the Bauer Core facility of Harvard University, and the Boston Children’s Hospital Molecular Genetics Core. This work was supported by the NIH (R01-GM123002 to LSC, F32-GM139244 to NJK) and the Ludwig Neurodegenerative Disease Seed Grants Program at Harvard Medical School. KC is supported by post-doctoral fellowships from the Fonds de Recherche du Québec - Santé and the Canadian Institutes of Health Research. BP and NK are supported by The Smith Family Foundation, NK is a Pew Scholar.

## Supplementary Information

## Supplementary Figures 1-9

**Table S1**. Gene level data of CRISPR screen hits.

**Table S2**. PREPL and NME6 immunoprecipitation mass spectrometry data.

**Supplementary Figure 1.**
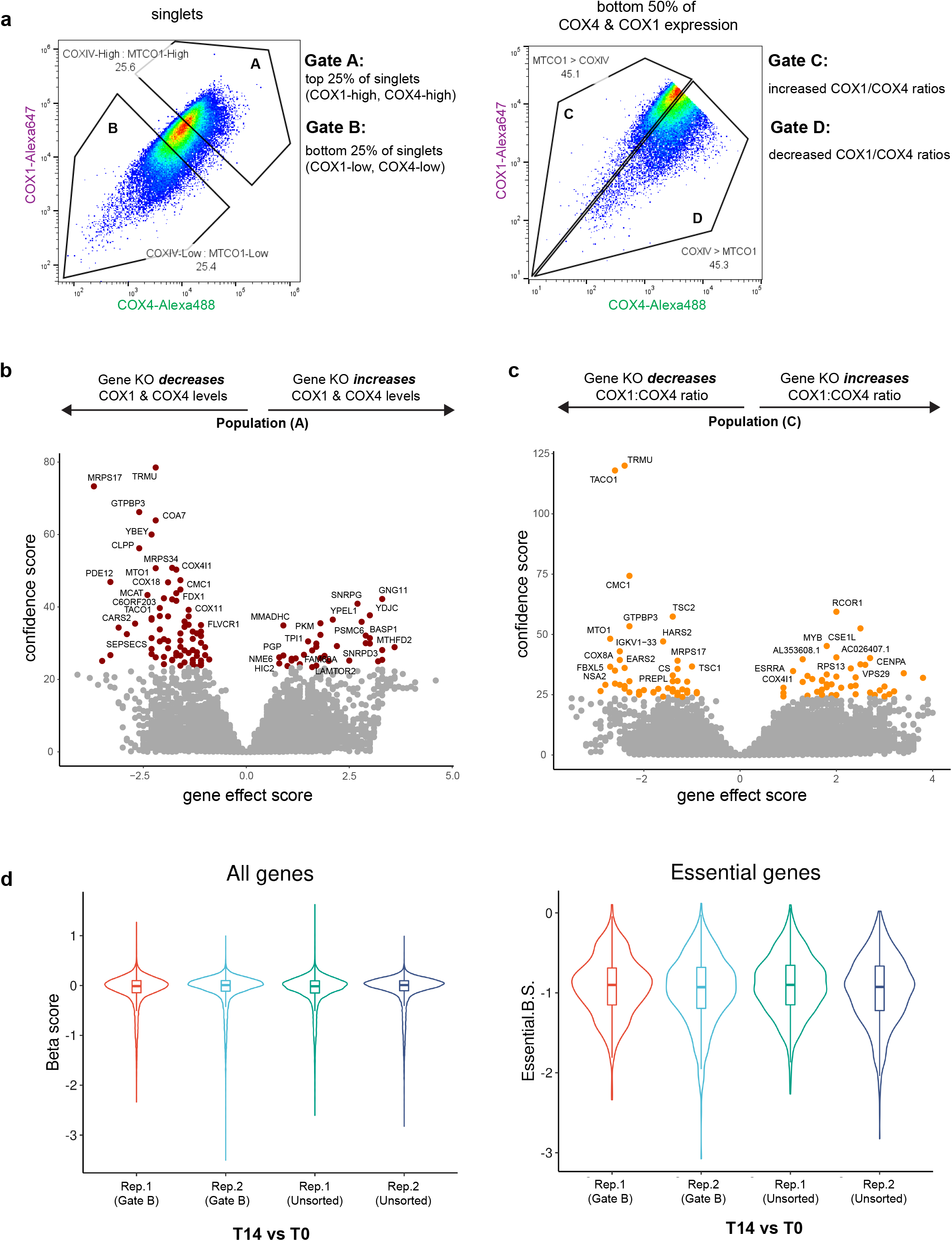
(**a**) Gating strategies used in flow cytometry depicting populations sorted for analysis in CRISPR screens. (**b**,**c**) Volcano plots of gene effect score (enrichment or depletion of sgRNAs in the sorted population relative to unsorted controls) vs confidence scores of genes identified in Population A and Population C for genes identified in ‘Gate C’ (COX1/COX4 ratio increased). Colored points = genes passing a 10% FDR threshold. (**d**) Quality control metrics for CRISPR screens showing violin plots of the beta scores (degree of selection, negative values = negative selection, positive values = positive selection; MAGeCK) for all genes (left) or known essential genes (right) for each sample after 14 days in culture (T14) relative to screen day 0 (T0). sgRNAs targeting known essential genes were negatively selected over time indicating functional Cas9 activity and consistent sgRNA library behavior.

**Supplementary Figure 2.**
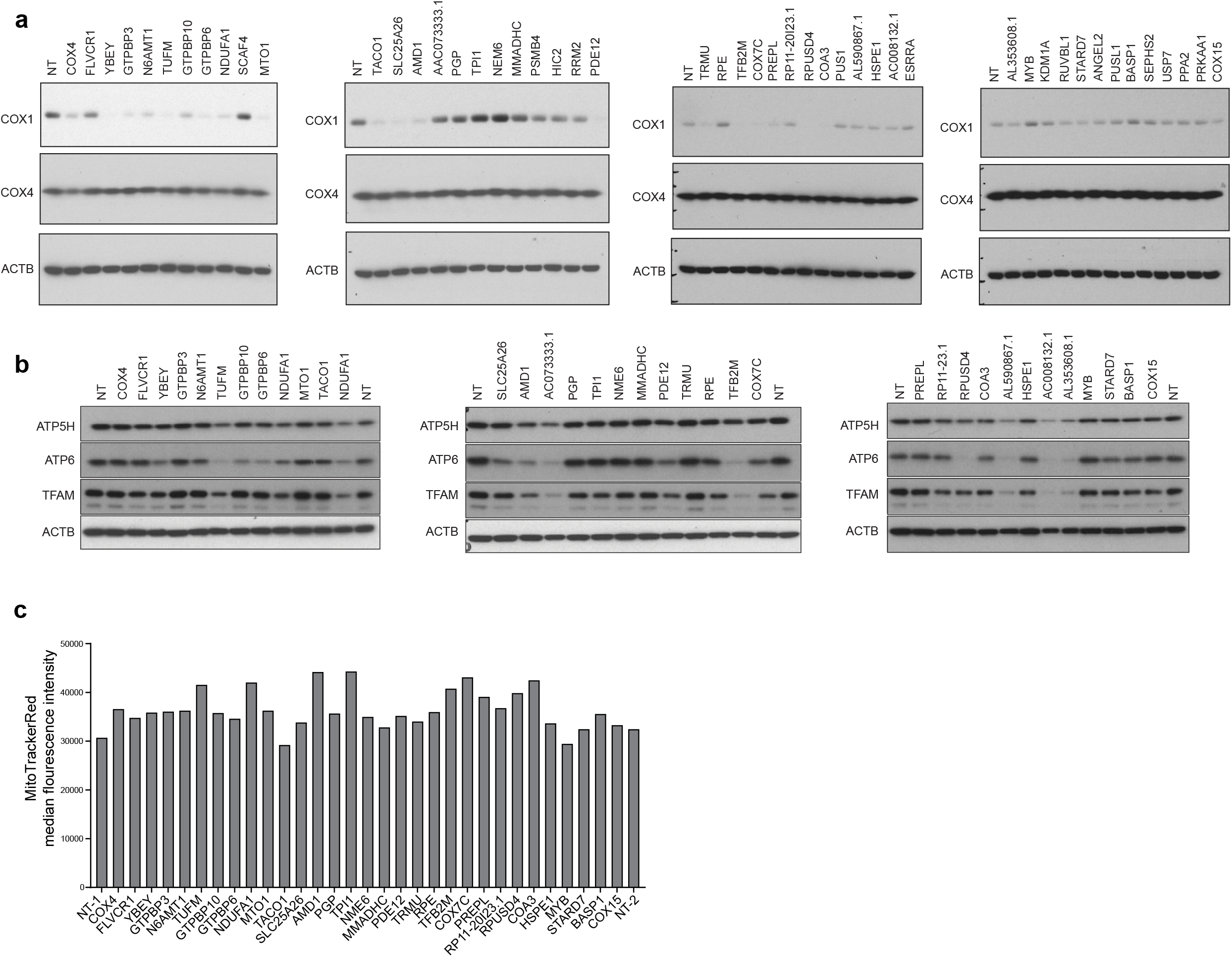
(**a**) Western blot validation of screen hits after individual sgRNA transductions (NT = non-targeting sgRNA). (**b**) TFAM and Complex V subunit levels (mito-encoded ATP6, nuclear-encoded ATP5H) levels in selected hits measured by western blotting. (**c**) FACS analysis of live cells transduced with individual sgRNAs and stained with MitoTracker Red CMXros (data are median fluorescence intensities from 20,000 analyzed cells).

**Supplementary Figure 3.**
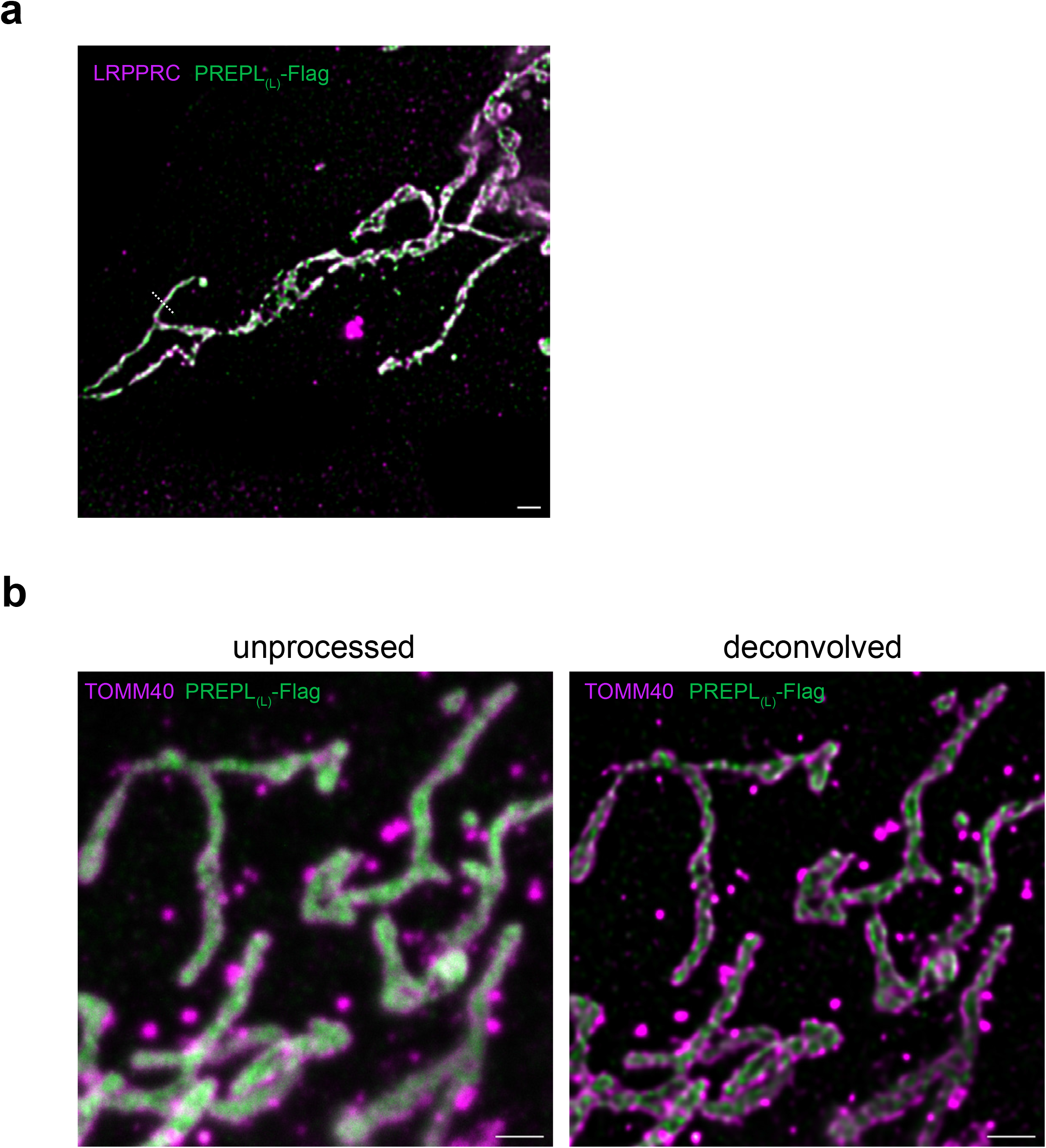
(**a**) Representative STED micrograph of LRPPRC (magenta; matrix marker) and PREPL_(L)_-Flag (green) co-staining. Quantification of line scans (across indicated dashed line) of pixel intensity for each fluorescence channel appear in **Fig. 3d**. (**b**) STED micrograph showing raw signal compared to deconvolved images using Hyugen’s deconvolution (default parameters, Scientific Volume Imaging; scale bars = 500 nm).

**Supplementary Figure 4.**
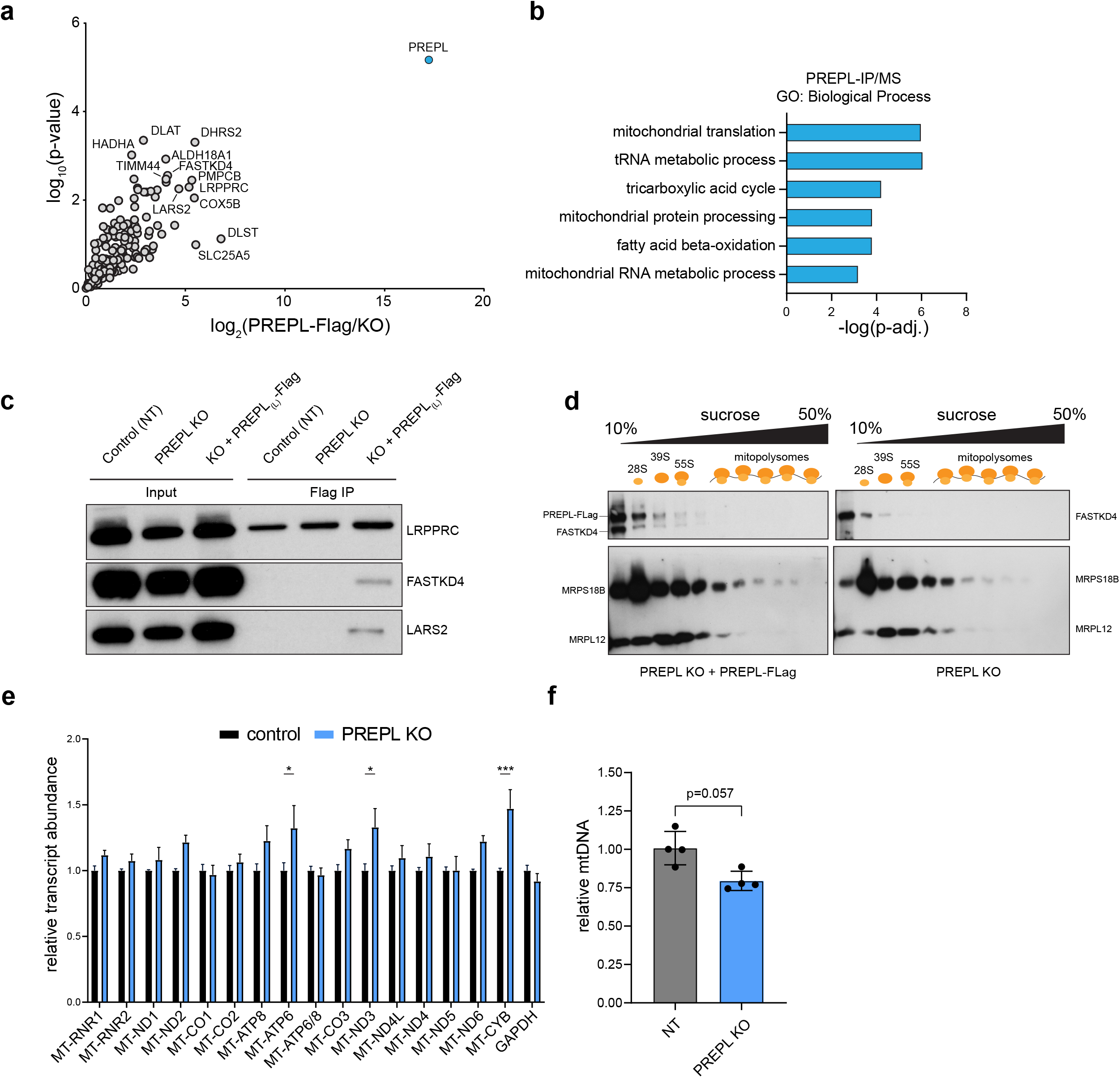
(**a**) Mass spectrometry identification of proteins in anti-Flag immunoprecipitations from PREPL KO or PREPL-KO + PREPL_(L)_-Flag whole cell lysates (PREPL_(L)_-Flag = bait; n = 3 replicates). (**b**) Enriched gene ontology terms for proteins identified as significantly enriched in PREPL_(L)_-Flag IPs relative to control KO IPs (Panther; Biological Process). (**c**) Co-immunoprecipitation (PREPL-Flag = bait) and western blot analysis of selected mass spectrometry-identified proteins. (**d**) Sucrose gradient centrifugation and fractionation with western blot detection of mitoribosome subunits, as well as PREPL_(L)_-Flag and FASTKD4 (MRPS18B = small mitoribosome subunit, MRPL12 = large mitoribosome subunits). (**e**) MitoStrings measurements of mt-RNA abundance in control and PREPL KO cells (n = 4 replicates from 2 independent experiments). (**f**) qPCR measurements of mtDNA levels normalized to nuclear DNA content (mito target = *MT-TL*, nuclear target = *B2M*; n = 4 replicates from 2 independent experiments, data are means +/- std deviation).

**Supplementary Figure 5.**
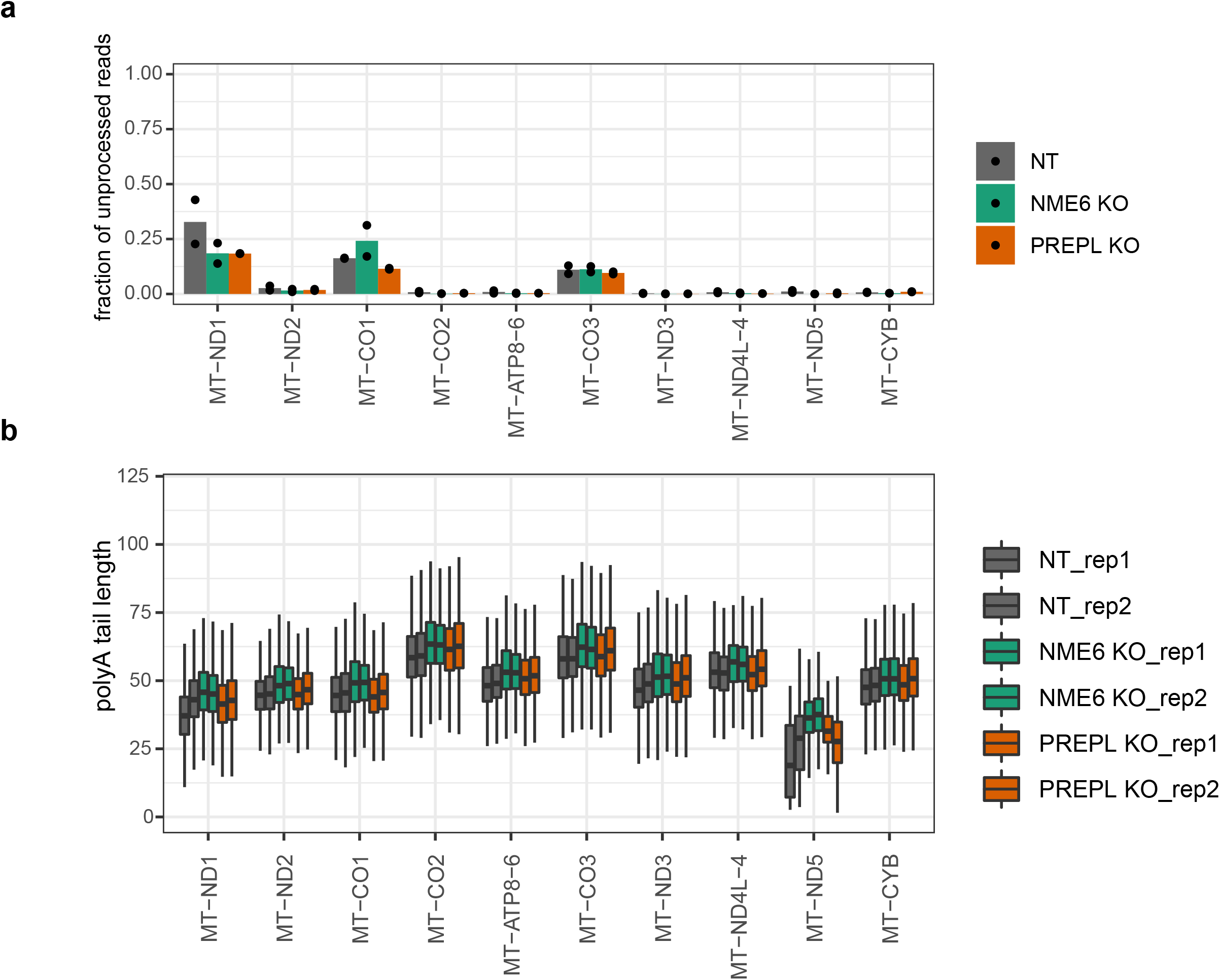
Direct RNA nanopore sequencing in control (NT), PREPL KO and NME6 KO cells. (**a**) Measurement of 5’-end processing levels for mt-mRNAs encoded on the heavy strand and (**b**) poly(A) tail lengths of mt-mRNAs in control, NME6 KO and PREPL KO cells. Two biological replicates are shown as dots in (a) and side-by-side in (b).

**Supplementary Figure 6.**
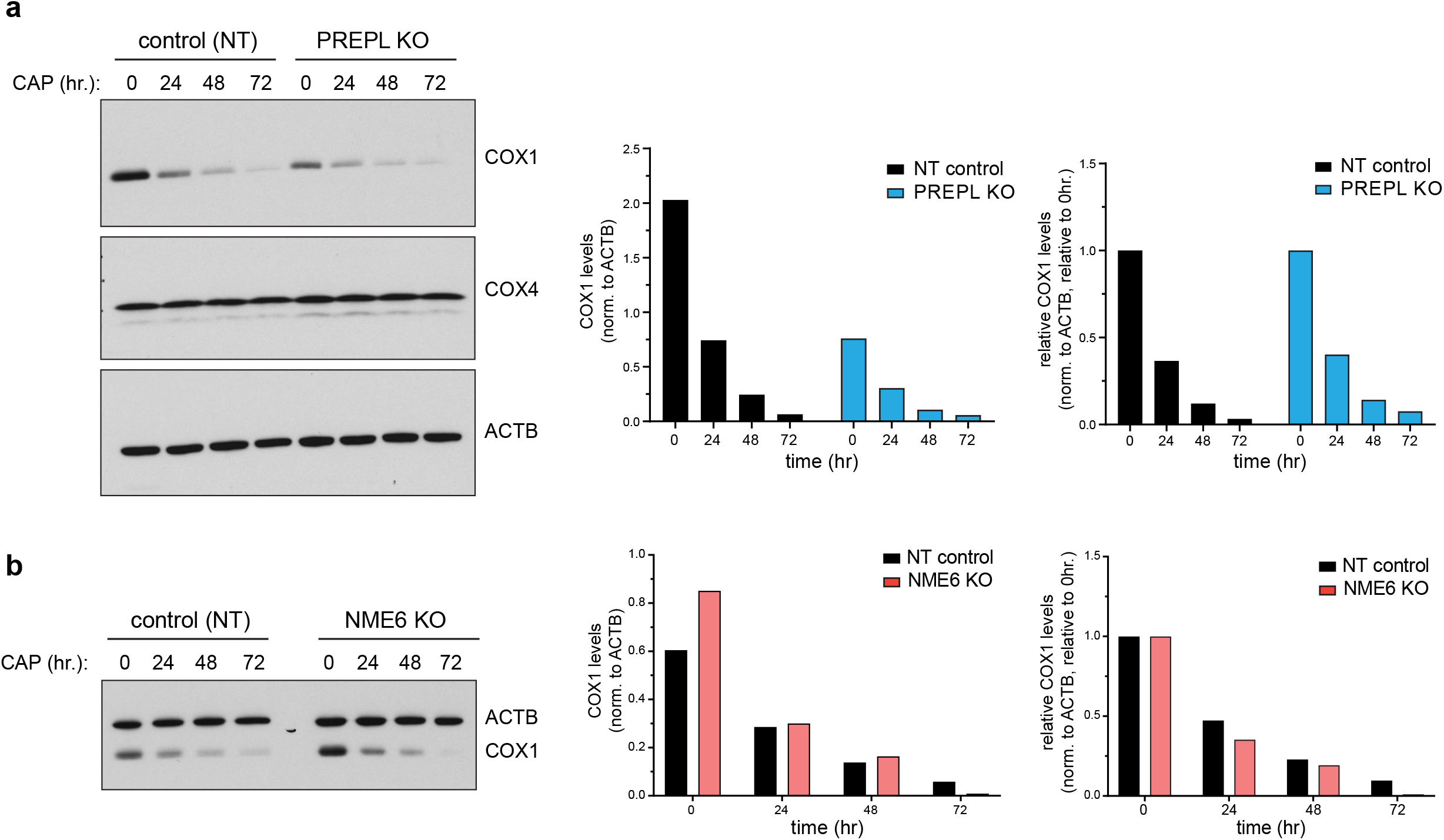
Mitochondrial protein turnover assays in (**a**) PREPL KO cells or (**b**) NME6 KO cells. Left panels: Cells were pulsed with 200 µg/mL chloramphenicol (CAP) to inhibit new mitochondrial protein synthesis for 0, 24 hr, 48 hr, or 72 hr., and proteins were detected via western blotting. Right panels: Quantifications of COX1 bands by densitometry were normalized to ACTB levels and plotted directly or normalized to time = 0 timepoints for each genotype to correct for the steady state subunit abundance levels in PREPL or NME6 KO cells.

**Supplementary Figure 7.**
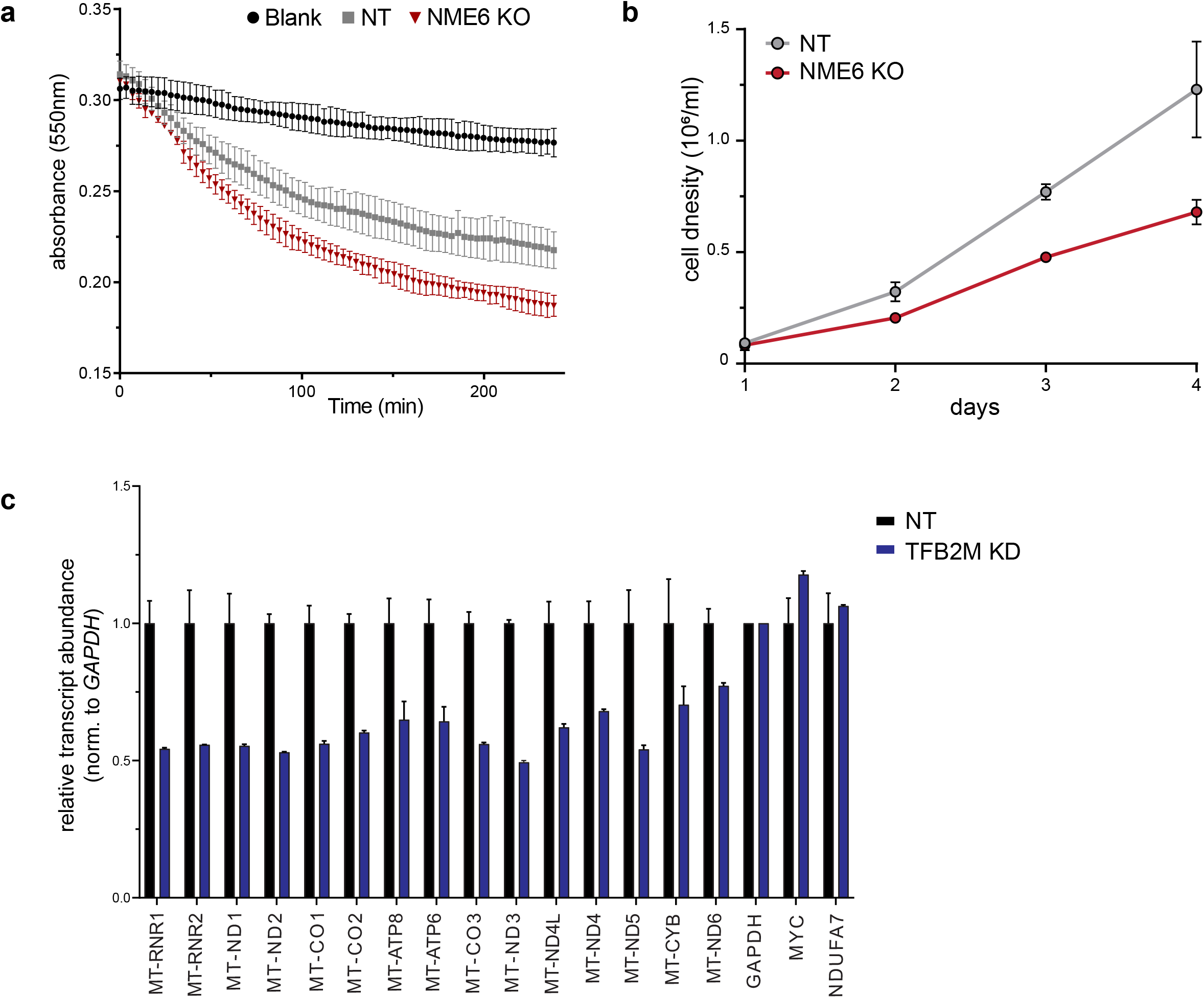
(**a**) Complex IV enzymatic activity measured colorimetrically by monitoring the oxidation of reduced cytochrome *c* over time from mitochondrial lysates immunocaptured on microplate wells coated with anti-Complex IV antibodies (related to **Fig 4i**). Shown here are the activities for 10 µg mitochondrial protein added per well in control and NME6 KO cells. Activities in OD/min were determined for **Fig 4i** by calculating the slope between 2 time points within the linear range of activity. (**b**) Proliferation of NME6 KO cells by hemocytometer counts of cell density at the specified days in RPMI-1640 media (n = 2 across independent KO clones, data are means +/- std deviation). (**c**) MitoStrings quantifications of RNA transcript abundance in control (NT) or *TFB2M* pooled CRISPR knock down cells (n = 2 independent cultures, data are means +/- std deviation).

**Supplementary Figure 8.**
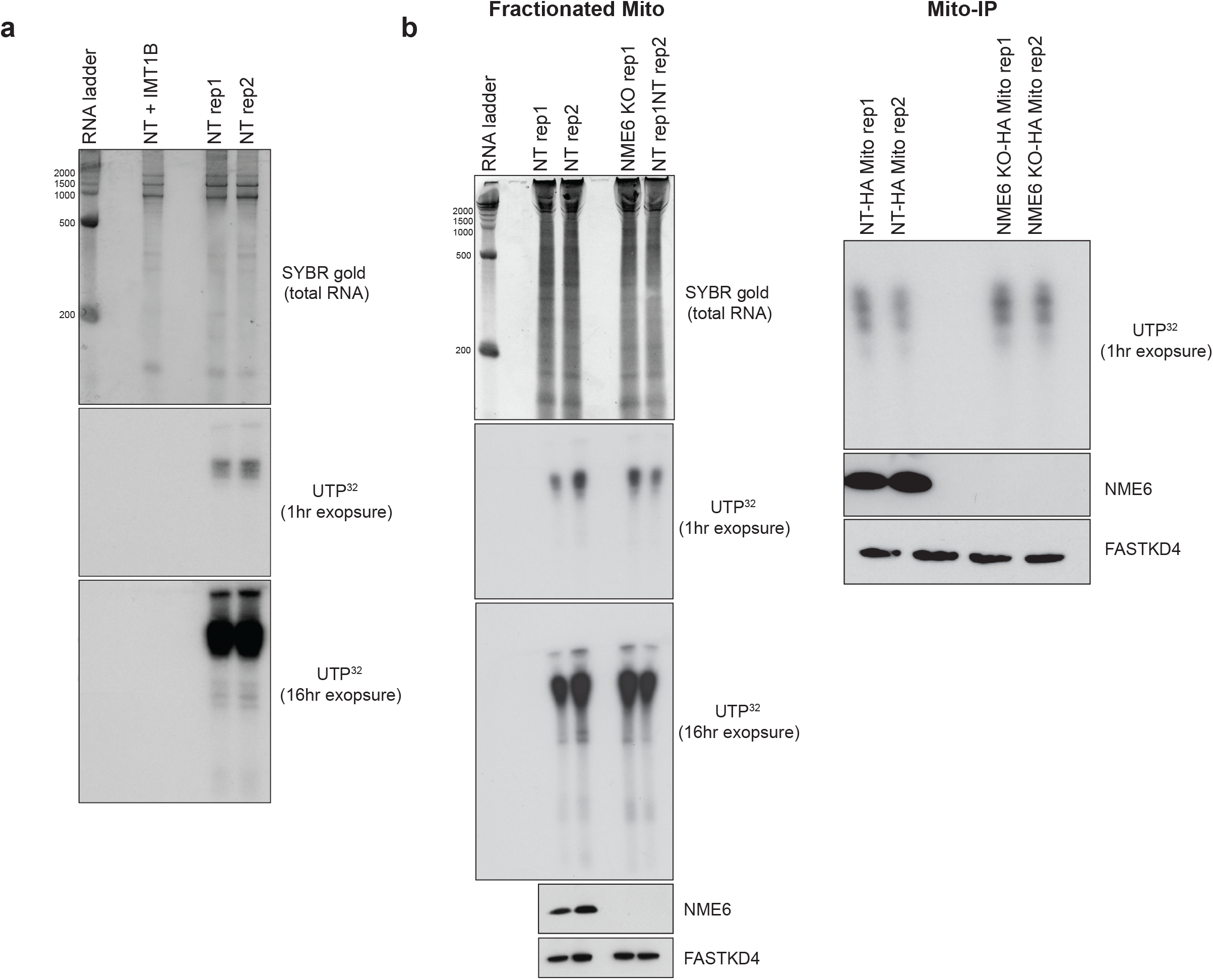
*In organello* mitochondrial transcription visualized by ^32^P-UTP labeling in purified mitochondria. RNA was visualized using TBE-urea PAGE, followed by autoradiography of newly synthesized mt-RNA. SYBR gold staining of total RNA and western blots on input mitochondrial lysates were included for controls. (**a**) IMT1B was used as a positive control to inhibit mitochondrial transcription in NT control cells (5uM, 2 hr. pre-treatment in culture and maintained in labeling reaction). (**b**) Transcription assay from fractionated or mito-IP purified mitochondria from NT and NME6 KO cells.

**Supplementary Figure 9.**
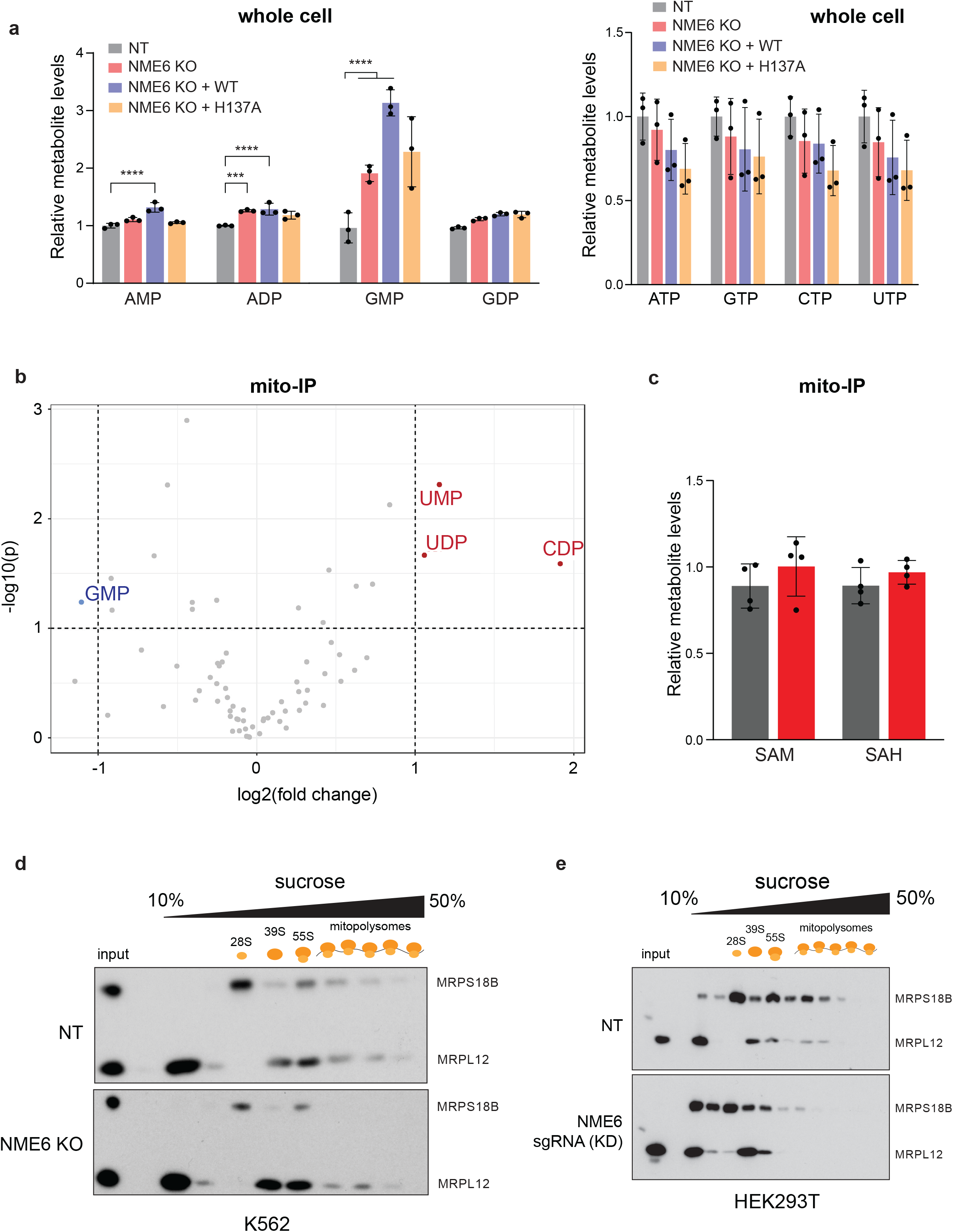
(**a**) Metabolite levels quantified by mass spectrometry in whole cell extractions from NT, NME6 KO, and WT/H137A rescue cell lines (relative to NT; *** p < 0.001, **** p < 0.0001; n = 3 independent cultures). (**b**) Volcano plot for confidently detected metabolites in mito-IP extracts comparing fold change in metabolites (NME6 KO/NT) vs significance values. (**d**) Metabolite levels in NT or NME6 KO mito-IP cell lines after HA-immunoprecipitation (relative to NT; n = 4 independent cultures). Bar plots are means +/- SD. SAM = S-adenosylmethionine, SAH = S-adenosylhomocysteine. (**d**) Sucrose gradient centrifugation and fractionation followed by western blotting of small (MRPS18B) and large (MRPL12) mitoribosome subunits to monitor mitoribosome assembly in (**d**) NME6 KO K562 cells and NME6 sgRNA-transduced pooled knockdown (KD) HEK293T cells (independent replicates of **Fig. 6e**).

## Materials and Methods

### Cell Culture

K562 cells were grown in RPMI-1640 media with GlutaMAX (Thermo Fisher Scientific), 10% FBS, penicillin (10,000 IU/mL), and streptomycin (10,000 ug/mL) unless otherwise noted. Cells used for CRISPR-Cas9 screens were cultured without penicillin/streptomycin, and were supplemented with uridine (50 ug/mL) and sodium pyruvate (1mM) to maintain mutant cells with dysfunctional mitochondria. Cells were maintained in a 37°C humidified incubator with 5% CO_2_. HEK293T and U2OS cells were cultured in DMEM with 10% FBS, penicillin (10,000 IU/mL), and streptomycin (10,000 ug/mL) in a 37°C humidified incubator with 5% CO_2_.

### Lentivirus packaging and transductions

Lentivirus were packaged in HEK293T cells following standard protocols for 3rd generation packaging. Lentivirus containing media was harvested 48 hours post-transfection of HEK293T cells and filtered using 0.45um PES membranes or centrifuged for 5 minutes at 300xg to remove cellular debris. K562 cells were transduced with lentiviral media using spin-infection methods. Filtered lentiviral supernatants were added to K562 cells with 8ug/mL polybrene and centrifuged for 2 hours at 1000xg, 33°C. Cells were resuspended in fresh media and expanded for 48 hours prior to selection. HEK293T or U2OS cells were transduced by adding viral supernatants directly to cell cultures (1:1 ratio viral supernatant to culture media) in the presence of 8ug/mL polybrene for 16-24 hr. then replaced with fresh media.

### Genome-wide CRISPR-Cas9 screens

K562 cells (ATCC) were transduced with lentiCas9-Blast (Addgene #52962) and single cell clones were selected for stable Cas9 expression. Clonal cell lines stably expressing Cas9 were used for CRISPR-Cas9 screening. Genome-wide CRISPR-Cas9 KO screens were performed as described in ^58,59^ with fixation and FACS staining as described in ^60^. Briefly, sgRNA libraries cloned into pMCB320 (Addgene #101926, #101927, #101928, #101929, #101930, #101931, #101932, #101933, #101934) targeting human protein coding genes (10 sgRNAs/gene) were packaged using 3rd generation lentivirus. 250×10^6 K562 cells were transduced with the genome-wide sgRNA library at an MOI < 1 and selected with puromycin (2ug/mL) 48 hr after transduction. A sample of cells was analyzed by FACS for mCherry expression to quantify transduction efficiency and successful sgRNA incorporation. Cells were expanded and the population was maintained 1000 fold over the number of sgRNAs elements to maintain library representation, while passaging cells every 2 days with fresh media. Cells in these screens were cultured without penicillin/streptomycin, and were further supplemented with uridine (50 ug/mL) and sodium pyruvate (1mM) to maintain any mutant cells with dysfunctional mitochondria. After specified time points (days 10, 12, and 14 post-transduction), cells were pelleted, washed with PBS, and fixed in ice-cold methanol (1mL per 10 million cells) for 10 minutes at -20°C (dropwise while vortexing) and then stained using standard intracellular FACS methods. Stained cells were sorted using a Sony SH800Z flow cytometer (∼30 million cells per gate) and immediately pelleted and lysed for genomic DNA extraction. Libraries were sequenced using an Illumina NextSeq platform and sgRNA compositions between unsorted control cells and sorted cells were compared using either casTLE^59^ and MAGeCK^61^ software packages.

### CRISPR KO clones

CRISPR mediated mutagenesis of target genes was performed by transducing K562 cells stably expressing Cas9 (lentiCas9-Blast, Addgene #52962) with lentivirus expressing sgRNAs (lentiGuide-Puro, Addgene #52963) targeting genes of interest. Transductions were performed as described above, and cells were selected with 2 ug/mL puromycin 48 hr after infection. Pooled mutant cells were then serially diluted to reach 1 cell/well in 96-well plates and grown up for ∼ 2 weeks. Single cell clones were screened by PCR and western blotting to confirm homozygous disruption of genes of interest.

### Immunoblotting and antibodies

Protein lysates used for immunoblotting were prepared in RIPA buffer in the presence of protease inhibitors, unless otherwise specified. Lysates were normalized using BCA assays, and standard SDS-PAGE and electroblotting protocols were used. The following antibodies were used in this study MT-CO1/COX1 (ABCAM, ab14705), COX4I1/COX4 (CST, 4850), ACTB (CST, 3700), FLAG (Millipore Sigma, F1804), LRPPRC (Abcam, Ab97505), TOMM40 (Proteintech, 18409-1-AP), MRPS18B (Proteintech, 16139-1-AP), MRPL12 (Proteintech, 14795-1-AP), WBSCR16/RCC1L (Proteintech, 13796-1-AP), NDUFS1 (Abcam ab169540), SDHA (Santa Cruz, sc-166947), UQCR4/CYC1 (Millipore Sigma, HPA001247), MT-ATP6 (ProteinTech 55313-1-AP), ATP5H (Abcam, ab110275), MT-ND1 (Abcam, ab181848), MT-CYB (ProteinTech 55090-1-AP), NME6 (Millipore Sigma, HPA017909), PREPL (Abcam, ab202064), CHCHD4 (ProteinTech 21090-1-AP).

### Subcellular mitochondrial and cytosolic fractionations

To obtain the cytosolic fractions, 5×10^6^ K562 cells were resuspended in PBS with EDTA-free protease inhibitor cocktail (Roche) and lysed using a 27.5G needle. Lysates were cleared by centrifugation for 10 min at 800x*g* at 4°C. The supernatant was collected and centrifuged for 10 min at 10,000x*g* at 4 °C to pellet organelles, and the supernatant was removed to obtain the cytosolic fraction. To obtain mitochondrial fractions, 30×10^6^ K562 were pelleted and resuspended in 800 µL hypotonic buffer (10 mM Tris, pH 7.5; 10 mM NaCl; 1.5 mM MgCl2) and left on ice for 7.5 min. Cells were then dounced in a 1mL glass homogenizer with 20 strokes on ice. 2M Sucrose T10E20 buffer (10 mM Tris, pH 7.6; 1 mM EDTA, pH 8.0; 2 M Sucrose) was added to the lysate to bring the sucrose concentration to 250 mM and homogenized cells were centrifuged at 600xg for 10 minutes to remove nuclei and unbroken cells. The supernatant was collected and spun at 10,000xg to pellet mitochondria. Pellets were then washed twice with 250 mM sucrose T10E20 buffer (10 mM Tris, pH 7.6; 1 mM EDTA, pH 8.0; 250mM Sucrose) before further application.

### Mitochondria immunoprecipitation

Mitochondrial immunoprecipitations were performed using K562 cells expressing an HA-mito tag (Addgene #83356) or a control MYC-mito tag (Addgene #83355) as previously described^37^. For immunoprecipitations, 30×10^6^ cells were pelleted and washed 2X in ice-cold PBS. Cells were then resuspended in 1mL PBS with 1× EDTA-free protease inhibitor cocktail (Roche) from which 5 μl of the cells were lysed in 50 μl of 1% Triton lysis buffer to obtain whole cell protein levels. The rest of the cells were lysed using 8 passages through a 30G needle. Lysates were spun for 1 min at 1000x*g* to pellet unbroken cells and supernatants were incubated with 50 μl HA magnetic beads(Thermo Fisher #88836) for 4 min.The beads were washed 3x in KPBS, and then lysed in 50 μl RIPA buffer for 10 min for western blotting or directly used in downstream applications.

### Proteinase K protection assay

20μg of isolated mitochondria, as described previously, (at a final concentration 1mg/mL in 250mM Sucrose T10E20 buffer) was either left untreated or was treated for 30 min on ice with 50μg/ml of proteinase K alone or in combination with 1% Triton-X100. After the proteinase K treatment, 2μl of 40 mM phenylmethanesulfonyl fluoride (PMSF) was added to all samples, and samples were incubated on ice for 20 min. The cells were then lysed in RIPA buffer with 1× EDTA-free protease inhibitor cocktail (Roche) and analyzed using western blots.

### RNA abundance measurements by nanoStrings

Total RNA was purified from cells using Trizol and isopropanol precipitations. 25-50 ng of total RNA per sample was used for each NanoStrings run. To hybridize RNA to the probe sets, first probe stocks ‘A’ and ‘B’ were diluted into working stocks by adding 4ul of each master stock into 29 ul TE + 0.1% Tween-20. Next, 70ul of nCounter Sprint hybridization buffer was added to a Nanostrings TagSet, then 7ul of the probe ‘A’ working stock, followed by 7ul of probe ‘B” working stock were each added to the TagSet. In a 0.2mL PCR tube, 8ul of TagSet/Probe mastermix + 25-50ng RNA/H20 was added for a 15ul total reaction volume. Samples were hybridized in a PCR machine at 67C for 16hr. before loading into an nCounter Sprint Cartridge for quantification.

### qPCR mtDNA levels

DNA was isolated from cells using a Qiagen Blood and Tissue DNeasy kit. Quantitative PCR was performed using Sso EvaGreen Supermix (BioRad), adding 60ng of template DNA per reaction and using mtDNA and nucDNA specific primers (400nM final concentration). *MT-TRNA-LEU* (mtDNA target): Fwd = CACCCAAGAACAGGGTTTGT, Rev = TGGCCATGGGTATGTTGTTA. *B2M* (nucDNA target): Fwd = TGCTGTCTCCATGTTTGATGTATCT, Rev = TCTCTGCTCCCCACCTCTAAGT. PCR was performed using a BioRad CFX384 Touch Real-Time PCR Detection System, with the following cycling conditions: 50°C for 2 min, 95°C for 10 min, 40 cycles [95°C for 15 sec, 62°C for 60 sec], + melt curve.

### STED microscopy

Stimulated emission depletion microscopy was performed on U2OS cells transduced with lentivirus expressing Flag-tagged PREPL cDNA (pCIG3, Addgene #78264). Cells were seeded 3 days post-transduction on coverslips (thickness #1.5H, Thor Labs cat# CG15NH1) and fixed in 4% paraformaldehyde for 20 minutes. Standard immunocytochemistry techniques were used to stain cells with the following antibodies (anti-FLAG M2 clone 1:200; anti-TOMM40 1:200 Proteintech cat# 18409-1-AP; anti-LRPPRC 1:200 Abcam cat# Ab97505). Secondary antibodies: goat anti-mouse Alexa-555 (1:100); goat anti-rabbit Alexa-647(1:100). Deconvolution was performed using Hyugen’s deconvolution (default parameters; Scientific Volume Imaging).

### Electron microscopy

K562 cells were pelleted and fixed in 2.5% Glutaraldehyde 1.25% Paraformaldehyde and 0.03% picric acid in 0.1 M sodium cacodylate buffer (pH 7.4) for 2 hr at RT. Samples were then processed by the HMS Electron Microscopy Facility as described: fixed cells were washed in 0.1M cacodylate buffer and postfixed with 1% Osmiumtetroxide (OsO4)/1.5% Potassiumferrocyanide (KFeCN6) for 1 hour, washed 2x in water, 1x Maleate buffer (MB) 1x and incubated in 1% uranyl acetate in MB for 1 hr followed by 2 washes in water and subsequent dehydration in grades of alcohol (10 min each; 50%, 70%, 90%, 2×10min 100%). The samples were then put in propyleneoxide for 1 hr and infiltrated ON in a 1:1 mixture of propyleneoxide and TAAB (TAAB Laboratories Equipment Ltd)). The following day the samples were embedded in TAAB Epon and polymerized at 60°C for 48 hrs. Ultrathin sections (about 60 nm) were cut on a Reichert Ultracut-S microtome, picked up on to copper grids stained with lead citrate and examined in a JEOL 1200EX Transmission electron microscope or a TecnaiG2 Spirit BioTWIN and images were recorded with an AMT 2k CCD camera.

### Complex IV enzymatic activity assays

Mitochondria were isolated from cells as described above using hypotonic lysis and differential centrifugation. Proteins were extracted from isolated mitochondria in the presence of protease inhibitors (Halt Protease Inhibitor Cocktail, Thermo Fisher) and protein content was quantified using BCA assays. Complex IV enzyme activity was measured colorimetrically in antibody-coated microplates by measuring the oxidation of cytochrome-c according to the manufacturer’s instructions (Abcam cat# ab109909).

### Pseudouridine Sequencing

CMC (N-cyclohexyl-N′-(2-morpholinoethyl)carbodiimide metho-p-toluenesulfonate; Santa Cruz Biotechnology) based detection of pseudouridine RNA modifications was performed as described in ^41,62^. Briefly, mitochondria were fractionated from cells as described above and RNA was purified using standard Trizol and isopropanol precipitation methods. 3ug of RNA from each sample was split into -CMC and +CMC conditions (2ug used for +CMC and 1ug used for -CMC) and adjusted to 20ul each with water. RNA was denatured by adding 2.9ul of 20mM EDTA pH8.0, heated for 3 min. at 80C, then returned to ice. 0.5M CMC was prepared fresh in BEU buffer (50 mM bicine pH 8.5, 4 mM EDTA, 7 M urea) just before use, and 100ul of either CMC or BEU buffer was added to -CMC or +CMC samples respectively and incubated at 40°C for 45 min. mixing at 1000 rpm in a thermomixer. RNA was precipitated (sodium acetate, 100% ethanol, glycoblue) and then CMC bound to U/G residues was reversed by heating RNA samples at 50°C for 2 hr. with mixing at 1000 rpm in sodium carbonate buffer (pH 10.4, 50 mM Na_2_CO_3_, 2 mM EDTA). RNA was precipitated (sodium acetate, 100% ethanol, glycoblue), washed, and resuspended in 15ul of water. RNA was quantified using a Nanodrop, and RNA-sequencing libraries were made using the SMARTer Stranded Total RNA Sample Prep Kit - HI Mammalian (Takara). Libraries were sequenced at the Bauer Core Facility (Harvard University) using 150bp paired-end reads using an Illumina NovaSeq.

### Seahorse assays

K562 cells were seeded (150,000 cells/well) by centrifugation onto poly-L-lysine coated 96-well Seahorse assay microplates and allowed to equilibrate in Seahorse XF RPMI Medium, pH 7.4 (Agilent, 103576-10) for 60 min, in a non-CO2 controlled incubator before assays were performed. Respiratory parameters were measured using a Seahorse XFe96 analyzer according to manufacturer’s protocols with four basal measurements followed by sequential injections of oligomycin (1 μM), FCCP (2 μM), and rotenone (0.5 μM) with antimycin A (0.5 μM), where three measurements were taken after each injection.

### 35-S Metabolic labeling

K562 cells were grown at 5×10^5^ cells/mL in FBS-supplemented RPMI media. For ^35^S metabolic labeling, 3×10^6^ cells were washed with 1x with PBS and resuspended in labeling media (RPMI without cystine, methionine, and cysteine, MP Biomedical 1646454, supplemented with dialyzed FBS and glutamine). Cells were equilibrated for 30min at 37°C and 5% CO_2_ and then treated with 100µg/mL of anisomycin for 10min. 200µCi/mL of EasyTag labeling mix (^35^S-cysteine/^35^S-methionine, Perkin Elmer CAT NEG772007MC) was added for 30min or 60min, upon which cells were washed with 2x ice-cold PBS and lysed in RIPA buffer with 1× EDTA-free protease inhibitor cocktail (Roche). Lysates were then separated on a 17.5% acrylamide gel for 6h at 150V and transferred onto a 0.22µM nitrocellulose membrane using semi-dry transfer. The signal was then imaged using an autoradiograph.

### 4sU metabolic labeling

4sU incorporation into K562 cells was determined by measuring the fractions of unlabeled mtRNA by NanoStrings after 4sU-labeling and depletion, as described in^63^. In short, K562 cells were labeled with 500 µM 4sU for 30 min and lysed in Trizol. RNA was purified using isopropanol precipitations with the addition of spike-ins using *in vitro* transcribed RNA: ERCC-000148, as an unlabeled control. RNA was denatured at 60°C for 10 min and biotinylated using 5 µg/mL biotin-MTS (Biotium) in 20% dimethylformamide (Sigma), 20mM Hepes pH 7.4 and 1 mM EDTA with incubation for 30 min. Free biotin was removed using phase-lock heavy gel tubes (5prime), and then the mixture of unlabeled RNA and biotinylated RNA was incubated with streptavidin beads from uMACS Streptavidin kits (Miltenyi Biotec) following manufacturer’s protocol. Post incubation, the RNA/bead mixture was loaded on a uMACS column and washed with 100mM Tris pH 7.5, 10mM EDTA, 1M NaCl, 0.1% Tween 20 to collect unlabeled RNA in the flow-through. Collected RNA was purified using miRNeasy kits (Qiagen) with DNAse treatments(Qiagen). MitoStrings measurements were performed as described previously.

### Blue Native PAGE

Blue Native polyacrylamide gel electrophoresis (BN-PAGE) was adapted from previously published protocols^64^. In short, 50 μg of isolated mitochondria, as described above, were resuspended in 20μL sample buffer cocktail (5 μL NativePAGE sample buffer 4X, 8 μL of 5% digitonin, and 7 μL of water). The suspension was left on ice to solubilize for 20 min, upon which the lysate was cleared with centrifugation at 20,000g for 10 min at 4°C. 2 μL of Coomassie G-250 was added to the supernatant and the proteins were separated using NativePAGE 3%-12% gradient gels. During electrophoresis, inner chambers were filled with dark blue cathode buffer (Bis-tris 50 mM, Tricine 50 mM and 0.02% Coomassie blue G250) and outside chambers were filled with running buffer (Bis-tris 50 mM). Gels were run for 30 min at 150V, upon which the dark blue running buffer was replaced with light blue running buffer (Bis-tris 50 mM, Tricine 50 mM and 0.001% Coomassie blue G250) and run for an additional 90 min at 250V. Proteins were transferred onto a PVDF membrane using wet transfer methods at 200mA for 90 min, fixed with 8% acetic acid, rinsed twice with water, and then air dried. Dried membranes were then activated in 100% methanol for 5 min, followed by methanol and water rinses to remove coomassie staining, and then blocked using 5% skim milk in TBST. OXPHOS complexes were analyzed by immunoblotting with indicated antibodies towards each complex.

### Sucrose gradient fractionations

50×10^6^ cells K562 cells were grown at 5×10^5^ cells/mL for sucrose gradient fractionations. Upon removing media, cells were rinsed with ice-cold PBS and resuspended in 600µL of mitoribosome lysis buffer (0.25% lauryl maltoside,50 mM NH_4_Cl, 20 mM MgCl_2_, 0.5 mM DTT, 10 mM Tris, pH 7.5, and 1× EDTA-free protease inhibitor cocktail (Roche)). Lysates were clarified by centrifugation at 10,000 RPM for 5 min. To isolate mitoribosomes, 450 µL of clarified lysate was added on to 12 mL of 10-50% linear sucrose gradient and centrifuged at 40,000 RPM for 3h at 4°C using an SW41Ti rotor. Gradients were fractionated into 800 µL fractions using a BioComp instrument and mitoribosomes were followed with western blotting using antibodies against MRPL12 and MRPS18B.

### Nanopore direct RNA sequencing

Total RNA was extracted using Trizol and isopropanol precipitation, followed by polyA+ enrichment of mRNAs using a Dynabeads mRNA Purification Kit (Thermo Fisher, 61006). 500 ng of polyA+ RNA was used for input into Oxford Nanopore Technologies Direct RNA sequencing library preparation kit (SQK-RNA002). Steps were performed following the manufacturer’s protocol except for the following changes: 1) RNA CS was not added to the initial ligation mix and instead replaced with nuclease-free water and 2) ligation of the RT adapter was extended from 10 to 15 minutes. The resulting libraries were sequenced on a MinION device (Oxford Nanopore Technologies) for up to 72 hours. Sequencing reads were basecalled live with MinKNOW (release 20.10.3 or later) and filtered for basecalling threshold > 7. U bases were substituted for T bases in the resulting reads, followed by alignment to the reference hg38 genome using minimap2^65^ with parameters -ax splice -uf -k14. Multi-mapping reads were included in downstream analyses. 5’-end processing status of each read mapping to mt-mRNAs was determined by mapping the location of the 5’-end of the read relative to the start of the transcript (−15 to +50 nt from transcript start). Reads that started in the transcript start window of a gene were classified as “processed” and reads that started upstream of this window were classified as “unprocessed”. Poly(A) tail lengths were measured using nanopolish ^66^. Reads with qc_tag “PASS” and with the 3’-end mapping within -15 to +15 nt of the transcript end were included in poly(A) tail length analyses.

### In organello mitochondrial transcription assays

Mitochondria were purified by subcellular fractionation or rapid mitochondrial immunoprecipitation as described above. Transcription assays were performed as described in^67^. Briefly, purified mitochondria were labeled with 30 μCi of alpha-32P UTP (Perkin Elmer BLU007H250UC) in transcription buffer (10 mM Tris pH 7.5, 25 mM sucrose, 75 mM sorbitol, 100 mM KCl, 10 mM K2HPO4, 50 μM EDTA, 5 mM MgCl2, 10 mM glutamate, 2.5 mM malate, 1 mg/ml BSA, 1 mM ADP) at 37°C for 20 min. then chased with cold UTP in transcription buffer for 5 min. at 37°C. Mitochondria were washed 3 times with wash buffer (10 mM Tris pH 6.8, 0.15 mM MgCl_2_, 10% glycerol) followed by RNA extraction using Trizol and isopropanol precipitation. RNA was resuspended in 2X urea sample buffer (Thermo Fisher) and separated by denaturing PAGE using Novex 6% TBE-urea gels (Thermo Fisher). Newly transcribed RNAs containing 32P-UTP were visualized by autoradiography.

### Mitochondria immunoprecipitation for metabolomics

Mitochondrial immunoprecipitation followed by metabolomics was adapted from^37^. Briefly,mitochondria were immunoprecipitated from 50×10^6^ K562 cells as described above. Pelleted cells were washed 2X with ice-cold KPBS, followed by homogenization using 25 strokes with a polytetrafluoroethylene (PTFE) [dounce homogenizer (VWR). Lysates were then centrifuged for 2 min at 1,000x*g* at 4°C. The supernatant was resuspended with 200 µL anti-HA magnetic beads and the mixture was incubated at 4 °C using an end-over-end rotator for 3.5 min. The beads were collected using a magnet to immobilize the mitochondria bound beads. Beads were washed using 3X ice-cold KPBS, and were immediately resuspended in either protein extraction or metabolite extraction buffer. Followed by metabolomics and western blot analysis as described in the methods.

### Metabolite Profiling by mass spectrometry for detection of polar metabolites

For whole cell measurements, 1×10^6^ K562 cells were washed with ice cold 0.9% NaCl briefly and metabolites were extracted using 200μL buffer (80% Methanol, 25 mM Ammonium Acetate and 2.5 mM Na-Ascorbate prepared in LC-MS water, supplemented with isotopically-labelled amino acid standards [Cambridge Isotope Laboratories, MSK-A2-1.2], aminopterin, and reduced glutathione standard [Cambridge Isotope Laboratories, CNLM-6245-10]) for 10min. The samples were vortexed for 10s, then centrifuged for 10min at 21,000g to pellet cell debris. The supernatant was dried on ice using a liquid nitrogen dryer. Metabolites were reconstituted in 20μL water supplemented with QReSS [Cambridge Isotope Laboratories, MSK-QRESS-KIT] and 1μL was injected into a ZIC-pHILIC 150 × 2.1mm (5μm particle size) column (EMD Millipore) operated on a Vanquish™ Flex UHPLC Systems (Thermo Fisher Scientific, San Jose, CA, USA). Chromatographic separation was achieved using the following conditions: buffer A was acetonitrile; buffer B was 20mM ammonium carbonate, 0.1% ammonium hydroxide in water. Gradient conditions were: linear gradient from 20% to 80% B; 20–20.5min: from 80% to 20% B; 20.5–28min: hold at 20% B at 150 mL/min flow rate. The column oven and autosampler tray were held at 25 °C and 4 °C, respectively. MS data acquisition was performed using a QExactive benchtop orbitrap mass spectrometer equipped with an Ion Max source and a HESI II probe (Thermo Fisher Scientific, San Jose, CA, USA) and was performed in positive and negative ionization mode in a range of m/z = 70–1000, with the resolution set at 70,000, the AGC target at 1 × 10^6^, and the maximum injection time (Max IT) at 20 msec. HESI settings were: Sheath gas flow rate: 35. Aux gas flow rate: 8. Sweep gas flow rate: 1. Spray voltage 3.5 (pos); 2.8 (neg). Capillary temperature: 300ºC. S-lens RF level: 60. Aux gas heater temp: 350ºC. An independent injection was performed for more specific detection of nucleotides. Mass spectrometer was operated in full scan mode in a range of m/z = 200–700 and m/z = 300–600 in negative and positive ionization mode respectively, with the resolution set at 70,000, the AGC target at 1 × 106, and the maximum injection time (Max IT) at 40 msec. HESI settings were: Sheath gas flow rate: 35. Aux gas flow rate: 8. Sweep gas flow rate: 1. Spray voltage 3.5 (pos); 2.8 (neg). Capillary temperature: 400ºC. S-lens RF level: 60. Aux gas heater temp: 450ºC.

### Data Analysis for metabolomics

Relative quantification of polar metabolites was performed with TraceFinder 5.1 (Thermo Fisher Scientific, Waltham, MA, USA) using a 7 ppm mass tolerance and referencing an in-house library of chemical standards. Pooled samples and fractional dilutions were prepared as quality controls and injected at the beginning and end of each run. In addition, pooled samples were interspersed throughout the run to control for technical drift in signal quality as well as to serve to assess the coefficient of variability (CV) for each metabolite. Data from TraceFinder was further consolidated and normalized with an in-house R script: (https://github.com/FrozenGas/KanarekLabTraceFinderRScripts/blob/main/MS_data_script_v2.4_20221018.R). Briefly, this script performs normalization and quality control steps: 1) extracts and combines the peak areas from TraceFinder output .csvs; 2) calculates and normalizes to an averaged factor from all mean-centered chromatographic peak areas of isotopically labeled amino acids internal standards within each sample; 3) filters out low-quality metabolites based on user inputted cut-offs calculated from pool reinjections and pool dilutions; 4) calculates and normalizes for biological material amounts based on the total integrated peak area values of high-confidence metabolites. In this study, the linear correlation between the dilution factor and the peak area cut-offs are set to RSQ>0.95 and the coefficient of variation (CV) < 30%. Finally, data were Log transformed and Pareto scaled within the MetaboAnalyst-based statistical analysis platform (PMID: 25897128) to generate PCA, PLSDA, and heatmaps. For mito-IP metabolomics, samples were normalized to total protein in purified mitochondria determined using BCA assays.

